# Pulsatile dynamics propagate crystalline order in the developing *Drosophil*a eye

**DOI:** 10.1101/2024.07.11.603179

**Authors:** Lydie Couturier, Juan Luna-Escalante, Khallil Mazouni, Claire Mestdagh, Minh-Son Phan, Jean-Yves Tinevez, François Schweisguth, Francis Corson

## Abstract

Pattern formation in developing tissues often involves self-organization guided by positional information. In most tissues, however, its dynamics, and therefore the underlying logic, remain unknown. Examining self-organized patterning of the fly eye, we combine experiments and modeling to elucidate how rows of light-receiving units emerge in the wake of a traveling differentiation front to form a crystal-like array. Live imaging of the proneural factor Atonal reveals unanticipated oscillations at the front, which are produced by the successive activation of two distinct enhancers and associated with pulsatile Notch signaling. Our observations are inconsistent with current models of eye patterning, whereby each row of differentiating cells provides a negative template for the next. Instead, they inform a new relay model in which transient Notch signaling from differentiating cells provides a positive template for the onset of differentiation two rows ahead, conveying both temporal and spatial information to propagate oscillations and crystal-like order.

## Introduction

There are many instances in development where a regular arrangement of cell fates self-organizes through cell-cell interactions; yet the dynamics by which these patterns arise, and the underlying logic, often remain elusive *(1)*. Lateral inhibition is a conserved self-organized patterning process in which Notch-mediated inhibitory signaling between equivalent cells results in the adoption of alternative cell fates *(2)*. It is commonly associated with disordered salt-and-pepper patterns. In the fly thorax, however, a broad prepattern of Notch receptor activation steers this self-organized system towards a stereotyped pattern of rows of sensory bristles *(3)*. Thus, self-organized Notch dynamics can be guided by a prepattern of Notch activity to produce a stereotyped outcome. Since Notch-mediated lateral inhibition occurs in many contexts in development *(4)*, we anticipated that the same logic, with different initial or boundary conditions, might capture the formation of diverse arrangements of cell fates. Here, we pursue this hypothesis by studying the logic of eye patterning in *Drosophila*.

### Ato expression oscillates in the eye

The adult eye exhibits a precise crystal-like array of ∼750 light-receiving multicellular units called ommatidia. Patterning starts with the formation of a regular lattice of R8 cells (Fig. 1A; each R8 is the founder cell of one ommatidia). This lattice forms over a 2.5 day-period in the eye imaginal disc as ∼30 rows of regularly-spaced R8s emerge sequentially behind a travelling differentiation front associated with a morphogenetic furrow (MF) *(5–7)*. Each R8 is selected by Delta-Notch signaling from groups of cells known as Intermediate Groups (IGs) showing high levels of Atonal (Ato), a proneural transcription factor required for R8 formation and eye development. Ato is expressed in the MF and in 3-4 rows of R8 cells (Fig. 1A) *(8–10)*. IGs form at regular spatial intervals along the posterior border of the proneural front. This pattern is thought to propagate via long-range activation and short-range inhibition. Hedgehog (Hh) produced by the differentiated R2 and R5 cells posterior to the MF activates Atonal expression in the MF, whereas Scabrous-regulated inhibitory Delta-Notch signaling from emerging R8s restricts the expression of Atonal to IGs *(5, 6, 11, 12)* (Fig. 1B). A pattern of staggered rows thus emerges as the MF progresses from posterior (P) to anterior (A) and each new row provides a negative template for the next. This negative templating logic naturally arises in a model for self-organized patterning by Notch *(3)* if we incorporate a receding inhibitory front that mimics the activity of the anterior inhibitors Hairy and Extramacrochaete (Emc) *(13)* (SI S7, Fig. S1, movie S1). Patterned cell rearrangements along the posterior margin of the MF may also contribute to R8 selection *(14)*. Importantly, however, all current models share the premise that spatial information is relayed between successive rows, from row *n* to *n+1 (11, 14, 15)*.

**Figure 1:**
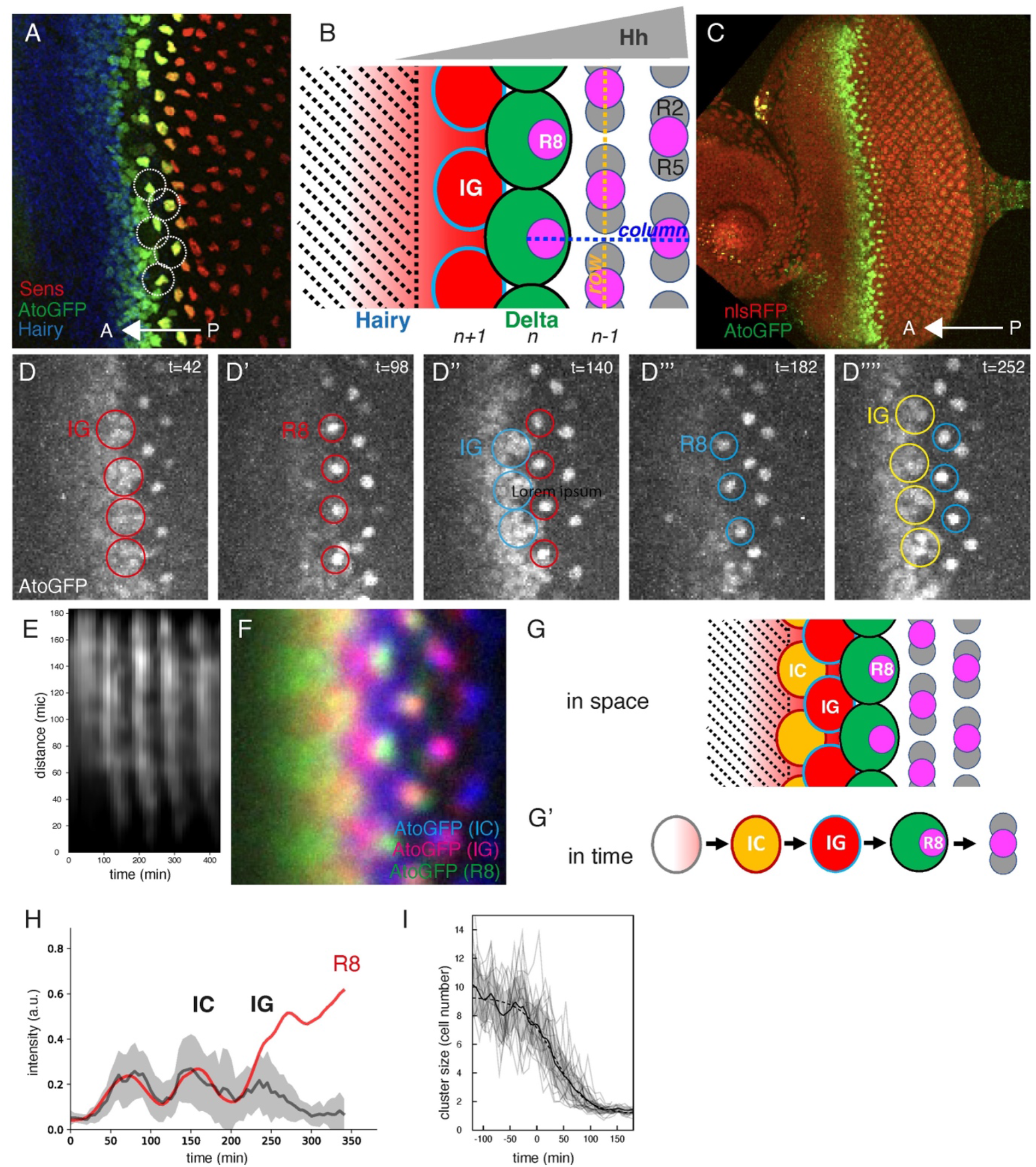
Oscillations of Atonal. (**A**) Undifferentiated cells (Hairy, blue) produce a regular pattern of R8s (Senseless, red) as they travel through a moving differentiation front (AtoGFP, green). (**B**) Patterning cues propagate from row *n* to row *n+1* via local inhibition of Ato by Delta and long-range activation by Hh. (**C**) Live imaging of AtoGFP (green; nlsRFP, red). (**D-D’’’’**) Each AtoGFP pulse produces a new row of IGs, which then resolve into R8s. (**E**) Kymograph of the AtoGFP signal along the MF, showing five Ato pulses. (**F**) Superposition of the registered pattern of AtoGFP at three stages showing that R8s (green) emerge posteriorly from IGs (red). (**G,G’**) Initial clusters (ICs) and intermediate groups (IGs). In space, ICs and IGs form two consecutive rows of staggered clusters (G); in time, the same group of cells undergoes an IC pulse followed by an IG pulse before an R8 cell is singled out (G’). (**H**) AtoGFP dynamics in one tracked R8 (red) and in its untracked neighbors (black). (**I**) Progressive resolution of the Ato clusters into R8s. The number of cells contributing to Ato clusters as a function of time (gray lines, 22 clusters from 2 movies; black line and gray region, mean±SD) is well fit by a sigmoidal function (dashed line; time scale,∼120 min). In all images, anterior (A; posterior, P) is left and time is in minutes (min).

To study patterning dynamics, we performed live imaging using GFP-tagged Ato (AtoGFP) in cultured eye discs. AtoGFP was detected at the MF, peaking in IGs and persisting in R8 cells over 3-4 rows (Fig. 1C), similar to endogenous Ato *(8, 10)*. Unexpectedly, however, pulses of AtoGFP were seen at the front (Fig. 1D-D’’’’, movie S2). Each new pulse coincided with the emergence of a new row of IGs, prefiguring a new row of R8 cells (the term ‘row’ refers to the alignments of IGs and R8s along the front; the term ‘column’ refers to the A-P-oriented succession of R8s, one in every other row; Fig. 1B). In our culture conditions, ∼3 pulses were routinely observed over a ∼6 hr period of imaging. Kymographs of the AtoGFP signal showed a period of 90 min +/-12 (n=8) (Fig. 1E, Fig. S2A-D), close to the rate of formation of new rows of ommatidia *(5)*. Live imaging also revealed that the pulses of Ato formed traveling waves along the front, running from the center of the disc towards the poles (movie S3). These traveling waves were detected in fixed samples in the form of localized pulses of endogenous *ato* mRNA and Ato protein expression (Fig. S2K-K’’). This implied that the *ato* gene undergoes pulses of transcription and that both Ato protein and *ato* mRNA are short-lived. Although oscillations were wholly unexpected, our observations are otherwise consistent with previous descriptions of Ato expression; in particular, a signature of traveling waves can be found in earlier reports of a gradient of IG maturation along the MF *(8, 10)*. To study the spatial and temporal structures of the Ato pulses, we applied a coarse-grained analysis to track the clusters of Ato-expressing cells. Using registered clusters to compute the average dynamics of Ato, we found that the pulse of Ato preceding R8 selection mapped to IGs and that R8s are selected from posterior IG cells (Fig. 1F, Fig. S2I-J, movie S4). This analysis also revealed that the preceding pulse mapped to groups of cells with elevated Ato expression, which had previously been identified as Initial Clusters (ICs) *(16)* (more anterior Ato expression, on the other hand, showed no clear spatial structure). In space, ICs and IGs form two consecutive rows of staggered clusters: ICs at row *n+2* belong to the same columns as emerging R8s at row *n*, whereas IGs at row *n+1* appear in alternating columns (Fig. 1G). In time, the same group of cells undergoes an IC pulse followed by an IG pulse before an R8 cell is singled out (Fig. 1G’). Consistent with this, single-cell tracking of randomly labelled cells *(17)* showed that MF cells undergo at least two Ato pulses (movie S5), and single-cell tracking of future R8 cells indicated that three pulses of AtoGFP can be seen in future R8 cells and in their untracked neighbors (Fig. 1H; n=3). We also quantified the weighted intensity of nuclear AtoGFP and the number of cells contributing to the tracked Ato clusters over time (Fig. S2E-H). The cluster size represented by this number decreased over time with a time scale of ∼120 min (Fig. 1I), showing that proneural cluster resolution is progressive. Thus, our analysis of Ato dynamics revealed that Ato undergoes several pulses in MF cells before becoming restricted to R8s.

### Ato oscillations are a composite of distinctly regulated pulses

Previous work has shown that two separate *cis*-acting enhancers regulate *ato* expression in the eye *(16)*. Initiation of *ato* expression and up-regulation in ICs involve the ato3’ enhancer, whereas expression in IGs and R8s is directed by the ato5’ enhancer. Our description of Ato dynamics suggested that the Ato pulse in ICs is directed by the ato3’ enhancer whereas the pulse in IGs is produced by the ato5’ enhancer (Fig. 2A-A’’). To test this, we generated transgenes that directed the expression of tagged-Ato in ICs (ato3’-AtoGFP) or IGs (ato5’-AtoHalo), as verified by FISH (Fig. 2B-B’’, Fig. S3D-E’). When combined, these two transgenes rescued the *ato* mutant phenotype (Fig. S3A-C) *(16)*. Live imaging showed that the ato3’ and ato5’ enhancers underwent synchronous pulses of activity (Fig. 2C-D, Fig. S3F-G, movie S6). Thus, spatially coherent oscillations at the front arise through the coincidence of distinctly regulated pulses at different positions. Considering individual cells over time, the succession of Ato pulses results from the composite addition of distinct pulses at different stages of specification. The basis of temporal periodicity in the expression of Ato in individual cells of the eye is thus analogous to the basis of spatial periodicity in the expression of pair-rule genes in the *Drosophila* embryo, where a periodic pattern is formed as the sum of independently regulated stripes *(18, 19)*.

**Figure 2:**
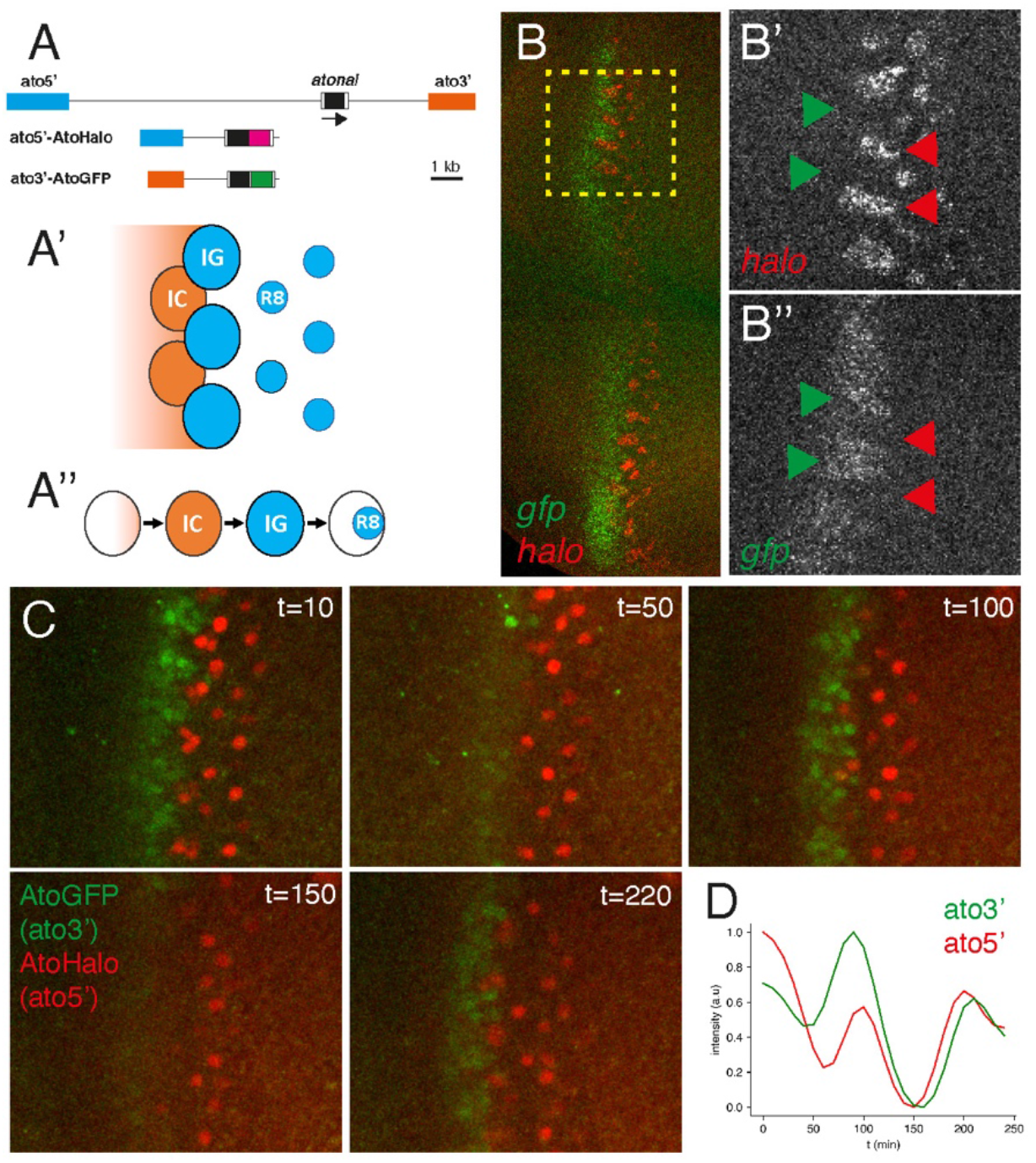
The composite addition of individual pulses produces oscillations. (**A-A’’**) Ato3’-AtoGFP and ato5’-AtoHalo reporters (A) show that coherent Ato expression across space (A’) and successive pulses in time (A’’) arise as the sum of 3’-and 5’-driven expression. (**B**-**B’’**) The ato3’ and ato5’ enhancers produce coordinated pulses of *ato* expression in ICs (green arrowheads; *gfp* probe, ato3’-AtoGFP) and IGs (red arrowheads; *halo* probe, ato5’-AtoHalo). (**C,D**) Distinct quasi-synchronous pulses are detected using ato3’-AtoGFP (green) and ato5’-AtoHalo (red). The GFP and Halo signals measured along the MF (from the kymograph in Fig. S3G) are plotted in D.

### E(spl) oscillates in both space and time

Since Notch is a key regulator of *ato* gene expression in the MF *(8, 10, 16, 20)*, we anticipated that Notch activity may likewise be dynamic in the MF. Examination of the expression of E(spl)-HLH Notch response factors indicated that E(spl)m*δ*-HLH (m*δ)* and E(spl)mγ-HLH (mγ) are expressed in the MF (Fig. S4). To study the dynamics of Notch signaling, we generated GFP-tagged versions of m*δ* and mγ, collectively noted as E(spl) in the following (Fig. S5). Live imaging revealed a dynamic pattern of teeth, arranged in successive rows with an alternating spatial phase (Fig. 3A-A’’, movie S7), producing a checkerboard pattern in kymographs (Fig. 3B, Fig. S6). Assigning a local phase to the pattern allowed us to register multiple positions and compute the average dynamics of E(spl) (Fig. 3C; Fig. S7A,D; movie S8). This showed that each new row of teeth emerged between the tips of the previous row, then extended anteriorly as the previous row subsided. Asking how this pattern relates to that of Ato, we first examined the patterns of the *ato* and *E(spl)* mRNAs *(8, 10)*. Based on individual samples (Fig. 3D-D’’), and a reconstruction of the dynamics from multiple samples (Fig. S8A, movie S9), we found that each new row of E(spl) teeth coincided with an Ato pulse, with IGs emerging between the teeth and ICs at their tips. Live imaging of GFPm*δ* together with a newly generated Halo-tagged Ato (AtoHalo) confirmed this conclusion (Fig. 3E-F; Fig. S7B,C,E; movie S10). Our observations, while consistent with earlier descriptions of Notch dynamics based on fixed samples *(10)*, reveal that a pulse of Ato in ICs maps in space with the tip of the E(spl) teeth, and coincides in time with a pulse of E(spl). Taken together, the correlation between the periodic patterns of Ato and E(spl) expression is consistent with a control of the dynamics by Notch. Yet the coincident expression and Ato and E(spl) in ICs ruled out a simple model in which spatially patterned expression of Ato is controlled by Notch-mediated inhibition.

**Figure 3:**
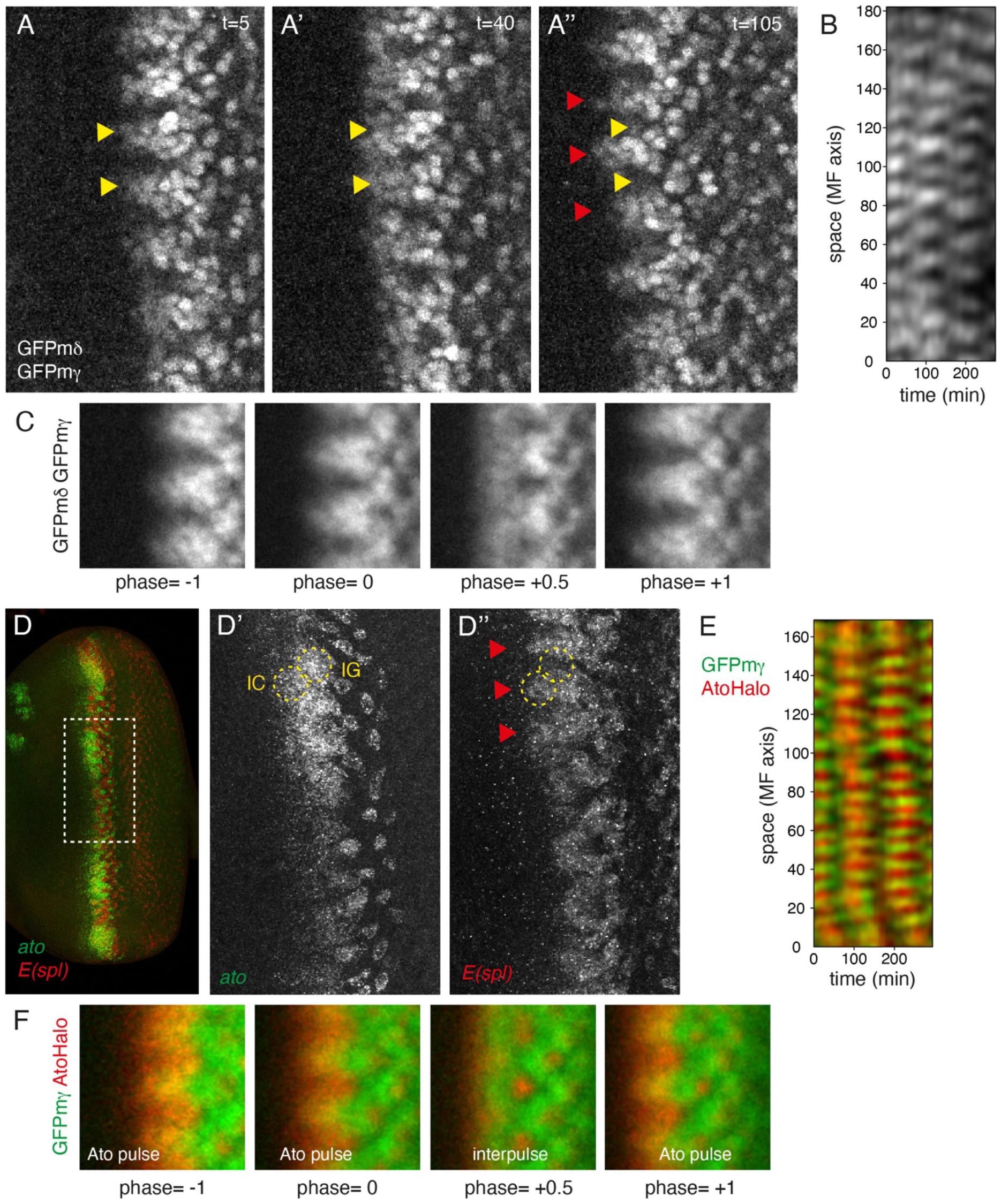
Periodic pattern of E(spl) (**A**-**A’’**) Live imaging of E(spl) reveals a periodic pattern of teeth (arrowheads) with an alternation between columns. (**B**) Kymograph of E(spl) along the front. (**C**) Average dynamics of phase-registered E(spl). (**D**-**D’’**) Ato pulses *(ato* mRNAs, green) co-occur with E(spl) teeth (arrowheads; *E(spl)* mRNAs, red). (**E**) Kymograph along the MF showing that Ato pulses (AtoHalo, red) coincide with the E(spl) teeth (GFPmγ, green). (**F**) Average dynamics of the phase-registered HaloAto and GFPmγ signals.

### A signal relay mechanism for pattern propagation

Our observations encouraged us to reconsider how spatial information is propagated in the eye. Whereas the emergence of IGs between E(spl) teeth is consistent with earlier models in which row *n* provides a negative template for row *n*+1, patterned Ato expression in ICs implied that spatial symmetry is broken as soon as row *n*+2, and its coincidence with the tips of E(spl) teeth suggested that Notch may instruct this early symmetry breaking. Since E(spl) teeth appear to emanate from the emerging R8s at their base, this suggested a model in which the pattern propagates through *positive* templating from emerging R8s in row *n* to ICs in row *n*+2 (Fig. 4A), rather than *negative* templating from R8s in row *n* to IGs in row *n*+1 as previously thought. Such a positive templating model, in which transient Notch signaling from emerging R8s provides both a spatial and a temporal cue for the onset of differentiation in ICs, also accounts for the recurrence of Ato pulses: as cells mature from IC to IG to R8, they in turn signal to induce a new cohort of more anterior IC cells in the same column (Fig. 4B). To account for both pattern propagation and R8 selection, this ‘relay model’ requires that Notch should transition from activating Ato in ICs to inhibiting Ato in IGs. Consistent with this, conditional inactivation of Notch resulted in the loss of the elevated, spatially patterned activity of the ato’3 enhancer in ICs, whereas in the posterior, proneural clusters failed to resolve and retained elevated Ato expression (Fig. S9A-D’’). On the other hand, a lower and more uniform ato3’ enhancer activity persisted in Hairy+ cells, as seen anterior to ICs in controls, reflecting the Notch-independent initiation of Ato expression at low levels in a pre-proneural state *(21)*. Notch may trigger elevated Ato expression in ICs directly, by relieving the transcriptional repression exerted by Su(H) (Fig. S9E-G) *(22)*, and perhaps also indirectly by down-regulating the inhibitor Hairy (Fig. S5F-G’)*(21)*.

**Figure 4:**
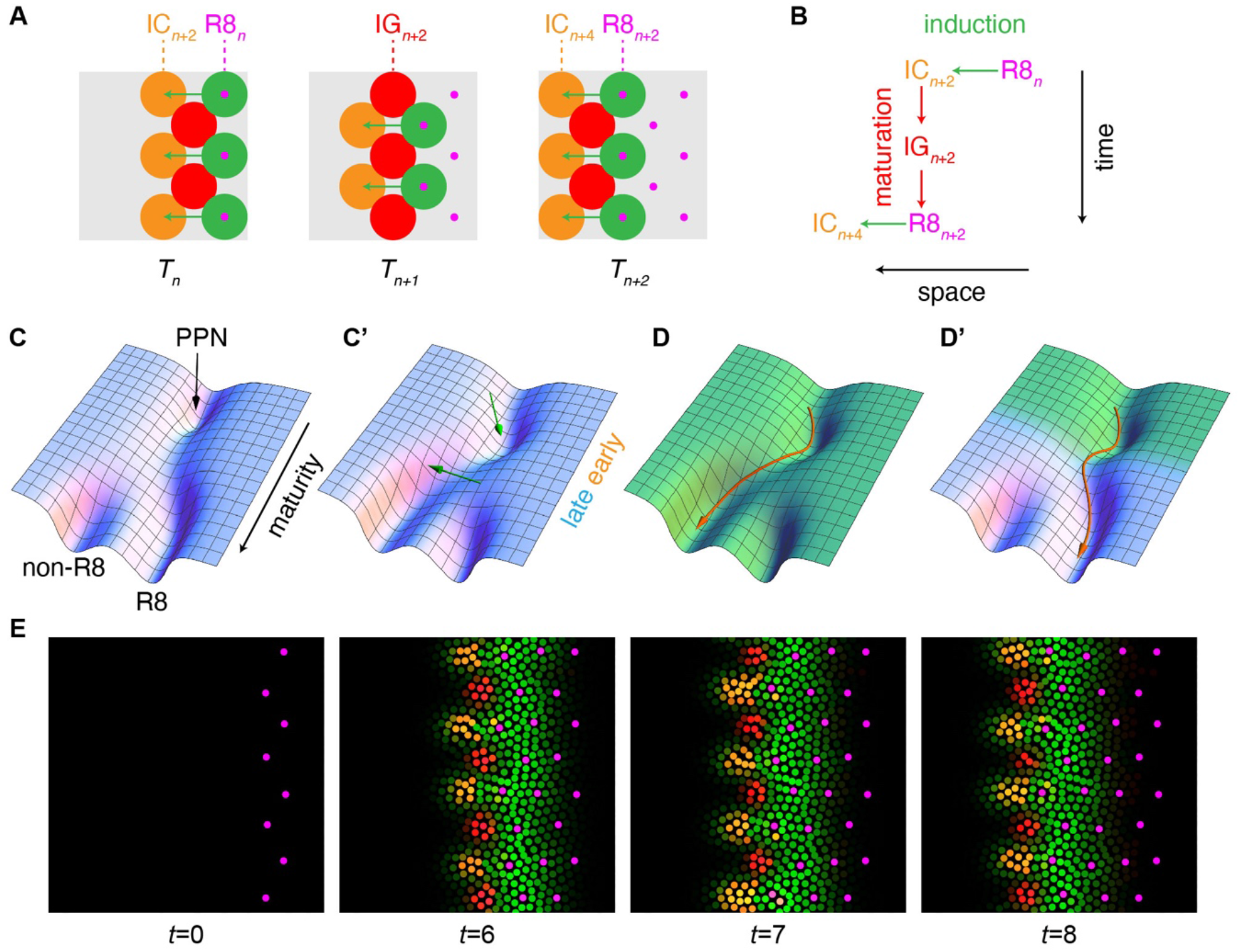
Relay model. (**A-B**) Sketch of relay model, in which a Notch-activating signal (green arrows in A) from late IGs/emerging R8s at row *n* induces differentiation in ICs at row *n*+2; these subsequently mature into IGs and R8s (red arrows in B), which in turn induce differentiation at row *n*+4 (**C,C’**) Landscape representation of dynamical model showing the response to different signal levels (C, low; C’, high). Notch activation is required for the exit from a pre-proneural state (PPN), after which maturing cells transition from an early regulatory regime in which Notch promotes proneural activity to a late, bistable regime in which Notch promotes a non-R8 fate (arrows in C’ represent the changing direction in which signaling ‘pushes’ cells in cell state space). (**D,D’**) Cell trajectories in response to different signaling histories. Under steady Notch signaling (represented by uniform green shading in D), proneural activity is initially promoted (cells move to the right) but cells eventually adopt a non-R8 fate. In response to a transient signal (reverting to a low level after cells have initiated differentiation, as represented by graded green shading in D’), cells instead adopt the R8 fate (to summarize the time-dependent landscape experienced by cells, cf. movie S11, the landscape in D’ interpolates between the high-signal and low-signal landscapes, cf. C’ and C, in the direction of increasing maturity). (**E**) Snapshots from a model simulation initialized with a regular template (green, signal *s*; reg/magenta, cell state *u*; cf SI S8).

Positive templating of ICs and negative templating of IGs could contribute redundantly to pattern propagation in the eye. On the other hand, the two models make different predictions for the response to conditional Notch inactivation: in a positive templating model (Fig. S10B,D), a memory of the patterned initiation of differentiation in ICs should persist, whereas in a negative templating model, in which Notch restricts uniformly activated Atonal expression, spatial information along the MF should be erased. Consistent with a positive templating model (Fig. S10A,C), *Notch* thermosensitive mutants at the restrictive temperature retain patterned expression of Ato in two rows of proneural clusters, which can be identified with the ICs and IGs that have been induced at the time of inactivation (Fig. S9A-D’’) *(12)*.

### A mathematical model recapitulates patterning dynamics

To test whether the proposed relay model can account for the dynamics of R8 patterning, we extended our mathematical model combining cell-intrinsic bi-stability and short-ranged inhibition *(3)* by incorporating an additional maturity variable. The logic of the resulting model, which builds in a signal-dependent onset of maturation and maturity-dependent dynamics (SI S8), can be summarized graphically in a landscape representation *(23*–*25)* (Fig. 4C,C’ and movie S11). Cell maturity starts increasing at a steady rate after cells have been pushed out of a pre-proneural state and roll down the landscape (Fig. S11A). Early in their maturation, cells respond to Notch by expressing Ato, whereas Ato dynamics in more mature cells is governed by the balance between self-activation and repression by Notch, giving rise to two distinct states - two distinct valleys - as in *(3)* (Fig. 4C’, Fig. S11B). Cells in which Notch is activated and stays on go to a non-R8 fate (Fig. 4D), whereas adoption of the R8 fate requires a signaling pulse followed by low Notch activity (Fig. 4D’). To capture the observed tooth-like pattern of signaling that supports long-range communication between rows, the range of signaling from cells at the late IG stage is allowed to transiently increase in the A-P direction. Finally, to build in a finite time window for fate specification, the dynamics of Ato is taken to become independent of signaling in sufficiently mature cells (Fig. S11C,D). Simulations of this model, initialized with a suitable template, captured the propagation of a periodic pattern of pulses, alternating between ICs and IGs within each A-P column and giving rise to a regular array of R8 cells (Fig. 4E, movie S11). Thus, modeling supports a relay model in which a Notch-activating signal that is largely autonomous to A-P columns mediates both pattern propagation from row *n* to *n+2* and the selection of R8 cells.

### 1D pattern propagation

The model predicts that the dynamics is largely autonomous to columns and should propagate in one dimension (1D) within an A-P column, going directly from row *n* to row *n*+2 in the absence of an intervening staggered row *n*+1 (Fig. 5A, Fig. S12A, movie S13). To test this, we studied the dynamics of patterning in clones of wild-type cells expressing AtoGFP and Halo-tagged mγ (Halomγ), surrounded by *ato* mutant cells. In thin A-P strips of wild-type tissue, the pattern propagated in 1D, and the temporal period of R8 emergence was twice that of Ato pulses, with non-R8-forming ICs alternating with R8-forming IGs (Fig. 5B-B’’, Fig. S12B-D). This 1D propagation is inconsistent with any model in which the initiation of Ato expression in ICs in row *n*+2 depends on spatial or temporal cues from row *n*+1 (Fig. S10G), as well as with a negative templating model (Fig. S10E; while this can also produce a 1D spacing pattern, its predicts that Ato expression in clusters of cells and R8 formation should recur with the same temporal frequency). Thus, the observed 1D propagation of the pattern strongly supports a relay model in which spatial and temporal cues are relayed from row *n* to *n+2*.

**Figure 5:**
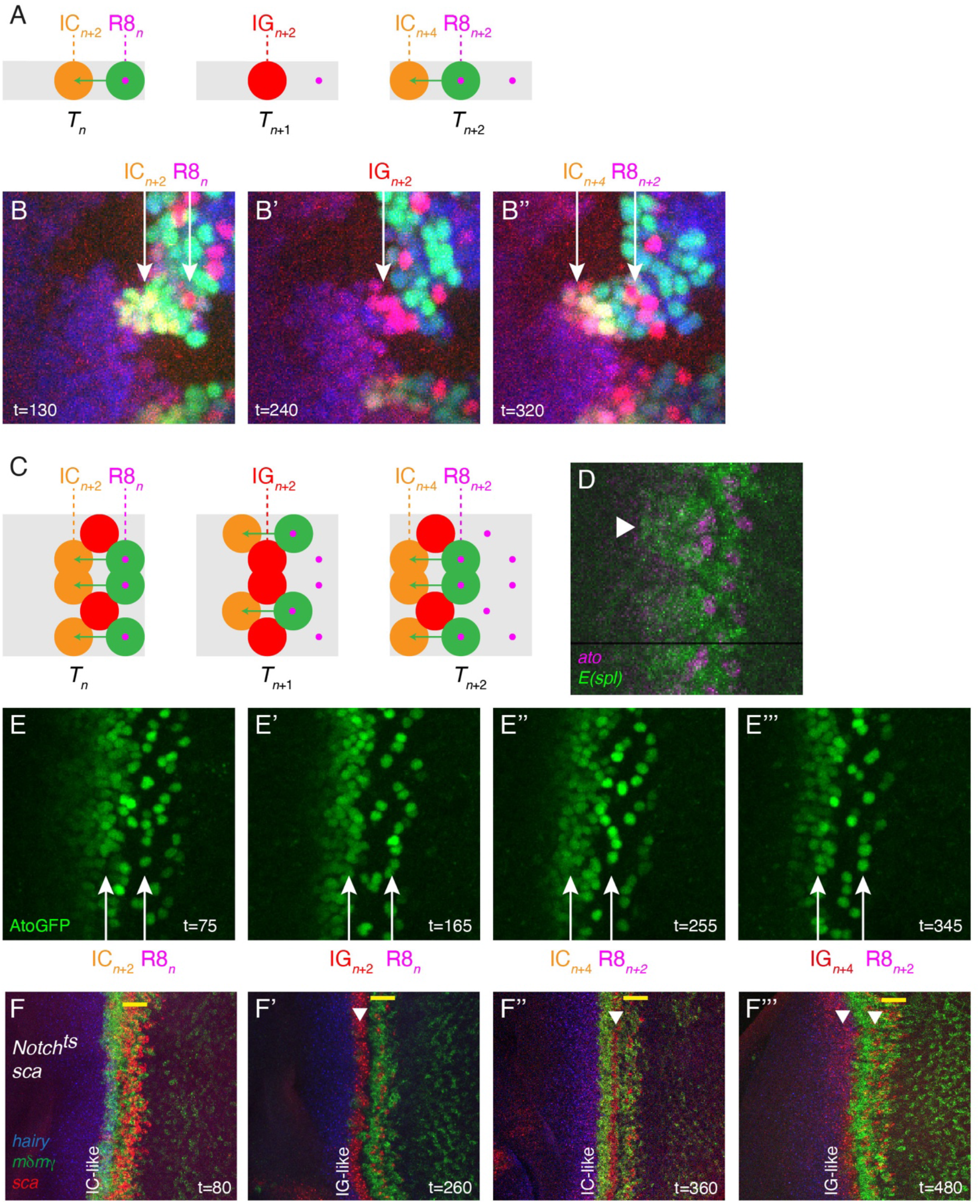
Altered pattern propagation. (**A**-**B’’**) 1D pattern propagation. Sketch of predicted dynamics (A; cf. also simulation in Fig. S13A and movie S13) and live imaging of *ato*^*1*^ mosaic disc (B-B’’) showing Ato (AtoGFP, red) and E(spl) (Halomγ, green) dynamics in a thin strip of wild-type cells at the MF (nlsRFP, blue; *ato* mutant cells unmarked; clusters are numbered n, *n*+2, *n*+4 with reference to the dynamics within a column in wild type, cf. Fig. 4A). (**C-E’’’**) Relaying of patterning defects. (C) Sketch of the predicted propagation of a defect (adjacent R8s in row *n)* to subsequent rows *(n*+2 and *n*+4). (D) Patterns of *ato* and *E(spl) (mδ, m*γ and *m8)* mRNA distribution in a *sca* mutant eye disk showing a broad E(spl) tooth anterior to adjacent R8s. (E-E’’’) snapshot from live imaging of AtoGFP in a sca mutant (cf. movie S14) showing the propagation of a defect (arrows denote adjacent R8s in rows *n* and *n*+2 and broad ICs and IGs in rows *n*+2 and *n*+4). (**F**-**F’’’**) Quasi-1D pattern propagation in 2D. A series of IC-like stripes *(E(spl)* mRNA, green; *hairy* mRNA, blue) maturing into IG-like/R8 stripes *(sca* mRNA red) were observed in *Notch*^*ts*^ *sca* double mutant growing at permissive temperature following a transient loss of *Notch* activity.

### Relaying of patterning defects

The relay model also has implications for the propagation of defects in the pattern, as in *scabrous (sca)* mutants. The *sca* gene encodes an endosomal glycoprotein interacting with Notch that is expressed in IGs downstream of Ato, and whose activity is required in R8s *(26*– *28)*. Loss of *sca* activity appears to reduce the range of lateral inhibition within IGs, leading to the frequent selection of more than one R8 cell per IG. The relay model predicts that when two or more R8s emerge from the same IG, signaling from these adjacent R8s should activate Ato in a broad IC, which after maturing into a broad IC will itself be prone to yield multiple R8s (Fig. 5C). As seen in simulations with a reduced baseline signaling range (Fig. S13B), the relay model thus predicts that defects should be relayed and repeated in the course of patterning. Consistent with these predictions, patterns of E(spl) and Ato in sca mutants showed broad E(spl) teeth and ICs anterior to groups of adjacent R8s (Fig. 5D), and live imaging of Ato GFP showed repeating cycles of broad IGs maturing into broad Ics yielding multiple R8s (Fig. 5E-E’’’, movie S14). Our model and observations thus identify an unexpected regularity in the disordered arrangement that develops in a patterning mutant.

### 1D in 2D

Whereas *sca* mutants exhibit an irregular pattern of IGs and R8s, experiments on *sca Notch*^*ts*^ double mutants have shown that this arrangement can be ‘reset’ through the transient inactivation of Notch, yielding a stably propagating pattern of near-continuous IG/R8 stripes. Whereas this pattern was previously taken to manifest a succession of proneural stripes that each resolve into a dense row of R8s *(15)*, the relay model predicted that - akin to a 1D pattern (cf. Fig. 5A,B) - it should propagate through the alternation of non-R8-forming IC-like stripes and R8-forming IG-like stripes. Time-course analysis of *sca* (used here as a marker of IGs and R8s *(27)*) and *E(spl)* expression validated this prediction: upon recovery from Notch inactivation, a stripe of IC-like cells formed and subsequently matured into a stripe of IG-like cells before resolving into a discontinuous stripe of R8s as a new IC-IG-R8 cycle began more anteriorly (Fig. 5F-F’’’). Consistent with these experiments, simulations with a reduced baseline signaling range showed that a uniform template could stably propagate for several cycles (Fig. S13E, movie S15) - an outcome that depended on the higher R8 density within rows: with wild-type parameters, by contrast, spatial and temporal order were spontaneously restored, at least locally (Fig. S13C,D; movie S16).

### A temporal bias in R8 selection

Beyond the propagation of an ordered array of ICs and IGs, simulations of our relay model also captured the experimentally observed posterior bias in R8 selection within IGs (Fig. S14). The model suggested that this spatial bias arises from differences in the timing of differentiation: posterior IC cells are exposed to Notch signaling earlier than more distal cells and, in the progression from IC to IG, are the first to initiate Ato self-activation and lateral inhibition. Analysis of the time courses of E(spl) expression in sparsely labeled cells *(17)* (Fig. S15, movie S17) supports this conclusion: a late onset of E(spl) expression (in anterior IC cells, identified as cells at the tip of E(spl) teeth) correlated with early re-expression of E(spl) at the IG stage, indicative of early exclusion from the R8 fate. Thus, a temporal bias may be sufficient to account for the observed spatial bias in R8 selection.

## Discussion

Our study of pattern formation in the *Drosophila* eye reveals the existence of Ato oscillations and pulsatile Notch signaling, informing a new relay model of R8 patterning. In this model, transient Delta signaling from emerging R8 cells at row *n* activates Notch in IC cells at row *n+2* to trigger their differentiation. Pattern is thus propagated by the relaying of spatial and temporal information across rows, such that spatial and temporal order are tightly intertwined. With pattern being propagated by positive templating from row *n* to row *n*+2, rather than by negative templating from row *n* to row *n*+1, our model reverses the previously accepted logic of pattern propagation in the eye; while negative templating could account for IG patterning, positive templating uniquely accounts for the existence of ICs, which had been observed previously *(16)* but remained uninterpreted. As in the patterning of sensory bristles in the fly thorax *(3)*, we find that Notch signaling underlies self-organized patterning through a dual role in the patterning of proneural clusters and the selection of neural precursors from among proneural cells. But whereas in the thorax (and in earlier negative templating models for eye patterning), both functions involve Notch acting as an inhibitor, in the eye Notch response transitions from activating (in the positive templating of ICs) to inhibiting (in R8 selection). This differential response is enabled by expression of Atonal under distinct enhancers, highlighting the evolvability of patterning by self-organized Notch dynamics that stems from the modularity of Notch activity outputs. The mathematical model we have introduced to describe the differential response of MF cells to Notch elaborates on earlier ‘geometric’ or ‘landscape’ models of fate specification *(23*–*25)*, by explicitly incorporating cell maturity as one dimension in the landscape down which cells ‘roll’ as they are being specified. In this representation, trajectories ending in either of two valleys capture the decoding of dynamical signals into fate.

Our relay model suggests mechanisms to evolve pair-rule patterns of ommatidia as observed in some insects *(29)* and has implications for the propagation and correction of errors in the pattern. Since the periodic emergence of ICs and IGs can proceed autonomously within A-P columns, the propagating pattern can be viewed as an 1D array of clocks, which must interact for adjacent columns to retain opposed phases and form R8 cells in alternation. While, in our simulations, Notch-mediated interactions are sufficient for this spatial and temporal order to emerge from a uniform template, other signals dynamically produced by patterned cells, such as EGF and Hh, could contribute to the tissue-wide coordination of patterning dynamics. A role for coordinated oscillations in developmental patterning has been extensively studied in the context of vertebrate somitogenesis *(30)*. In the eye as in the presomitic mesoderm, modulations in space of the natural frequency of an array of coupled clocks could account for the appearance of traveling waves. But the two systems differ in essential ways. Whereas in the somitogenesis clock, oscillations are understood to arise from cell-autonomous delayed negative feedback, in the eye Ato pulses recur because differentiating cells as they mature signal to more anterior cells to initiate differentiation, in what could be described as a ‘domino model’ for temporal periodicity. It is because the fundamental period of the dynamics is twice the period of Ato pulses, with a full cycle going from IC to IG and back to IC within a column, that adjacent columns can form R8 cells in alternation even though they undergo synchronous Ato pulses, supporting the propagation of a 2D pattern.

Our findings suggest an unusual route for the establishment of crystalline order. Whereas the addition of successive rows of R8 cells, each providing a template for the next *(6, 7)*, can be likened with the addition of successive layers in crystal growth, the patterning mechanism we describe, which proceeds through pulsatile signaling and gene expression, relies on the unique ability of living cells endowed with memory to encode and decode dynamical signals. We propose that in the eye disk, periodic pulses of Ato traveling from the center to the poles guide the orderly addition of new rows of R8s, building a tissue-wide spatial order that prefigures the nearly defect-free structure of the adult eye.

## Acknowledgements

We thank B. Hassan, D. Ish-Horowics, Y.-N. Jan, A. Jarman, D. Marenda, and F. Pignoni for reagents; S. Herbert for assistance with image analysis; and L. Bally-Cuif, V. Hakim, and E. Siggia for discussion and critical reading of the manuscript. We gratefully acknowledge UtechS Photonic BioImaging (Imagopole), C2RT, Institut Pasteur, supported by Agence Nationale pour la Recherche (ANR-10–INBS–04).

## Supplementary Materials

Materials and Methods

Figs. S1 to S15

Tables S1 and S2

Movies S1 to S17

