## Supplementary Information for "Pulsatile dynamics propagate crystalline order in the developing *Drosophil*a eye"

**This PDF file includes:**

Materials and Methods
Figs. S1 to S15
Tables S1 and S2
Captions for Movies S1 to S17
References

**Other Supplementary Materials for this manuscript include the following:**

Movies S1 to S17

**Materials and Methods**

### S1. Flies

The following transgenes and mutations were used: *ato*$^{1}$ (*9*), *ato*$^{w}$ (*31*), *ato*$^{GFP}$ (*32*), *N*$^{ts}$ (BL-2533), *sca*$^{1}$, *sca*$^{BP2}$, *ato3'(F5.8)-lacZ* (*16*), *ey-FLP* (BL-5576), *FRT82B ubi-mRFPnls* (BL-30555), *vasa-phiC31* (BL-40161), *vasa-Cas9* (BL-55821), *hs-Cre* (BL-851), Cad-3xmKate3 (*33*). Clones of *ato*$^{1}$ and *ato*$^{6Agfp}$ mutant cells were produced using the FLP-FRT system using *ey-FLP*. Trans-heterozygous *sca*$^{1}$/*sca*$^{BP2}$ mutants were studied here. *Notch* activity in *N*$^{ts}$ mutants was inhibited at 31°C for 2hr (Fig. 5F-F'''). Sparse labeling was achieved using a pUbi-FRT-stop-FRT-nlsHalo transgene as described in (*17*). Adult flies were imaged using a Zeiss Discovery V20 stereo-macroscope using a 1.0X (PlanApo S FWD 60mm) objective.

### S2. Genome engineering

The AtoHalo allele was produced by Recombination-Mediated Chromosomal Exchange (RMCE) at the *ato* locus (*31*). The donor plasmid for exchange was produced using multi-plasmid recombineering in *E. coli*. The Halo7 DNA fragment was obtained from UAS-Halo7 flies (BL-67619) using genomic PCR. Following sequencing, plasmids were injected in *vasa-phiC31*; *ato*$^{w}$ / TM6B *Tb* embryos and RMCE-positive chromosomes were selected based on loss of the *w*$^{+}$ marker. Correct recombination at the attP sites was verified by genomic PCR and sequencing. The resulting *ato*$^{Halo}$ allele appeared to be fully functional. Cloning details of donor and exchange plasmids are available upon request.

The *E(spl)m$\delta$-HLH (m$\delta$) and E(spl)m$\gamma$-HLH (m$\gamma$)* genes were individually GFP-tagged using Cluster Regularly Interspaced Short Palindromic Repeat (CRISPR)-mediated Homologous Recombination (HR). To do so, Cas9-expressing embryos were injected with a mix of plasmids encoding gRNAs flanking the region to be recombined together with a donor template carrying a 3xP3-RFP selection marker flanked by loxP sites. The following gRNAs were used:

*GFPm$\delta$*, 327 nucleotides (nt) 5' and 1219 nt 3' to the inserted GFP sequence, respectively:

gRNA1: GTGCAAGAAGAGAGCCTCAG

gRNA2: TGGATGGGTGGTGCATTACA

*GFPm$\gamma$*, 82 nt 5' and 958 nt 3' to the inserted GFP sequence, respectively:

gRNA1: GTTGTGTGTTGCTAGACCTT

gRNA2: GTAACTATTCGATCGCCCCA

Oligonucleotides were cloned into pU6-BbsI-chiRNA (Addgene #45946). Donor templates for HR were first produced by BAC recombineering in *E. coli* and then transferred into multicopy vector (*34*). Cloning details of donor templates will be provided upon request. The 3xP3-RFP selection marker was removed using Cre recombinase-mediated excision with an hs-Cre.
The *GFPm*$\delta$ *GFPm*$\gamma$ double-tagged chromosome was obtained in two steps. First, we used CRISPR-mediated HR to replace the *m*$\delta$ and *m*$\gamma$ genes by a *3xP3-miniwhite* ($w^+$) marker flanked by two attP sites using the following gRNAs:
gRNA1: GTGCAAGAAGAGAGCCTCAG
gRNA2: GTAACTATTCGATCGCCCCA
This modification was introduced in the *m7/8*$^{FRT}$ chromosome (*35*) such that the resulting *m*$\delta$*m*$\gamma^{attP,\ w+}$ chromosome also carried FRT sites flanking the *m7* and *m8* genes. Second, we used RMCE to exchange the *w+* marker with a *GFPm*$\delta$ *GFPm*$\gamma$ exchange plasmid obtained by multi-plasmid recombineering in *E. coli*. Injection was performed in *vasa-phiC31*; *m*$\delta$*m*$\gamma^{attP,\ w+}$ / TM6B *Tb* embryos. RMCE-positive chromosomes were selected based on loss of the $w^+$ marker and correct recombination was verified by genomic PCR and sequencing.

The *ato3'-AtoGFP*, *ato5'-AtoHalo*, *ato3'-nlsGFP* and *ato3'*$\Delta$*SBS-nlsGFP* transgenes were produced using SLIC and/or multi-plasmid recombineering in *E. coli*. The *ato5'* enhancer corresponds to the *ato5'-Eye* enhancer reported earlier (*16*). The *ato3'* enhancer corresponds to a 1.8 kb fragment encompassing the *ato_1.2* enhancer (*36*). This 1.8 kb fragment was defined such as to include the five SBS detected in this region (Fig. S9C). This 1.8kb fragment displayed similar rescue activity as the larger ato3'(F5.8) fragment (*16*) (Fig. S3). Cloning details will be provided upon request. The *ato3'-nlsGFP* and *ato3'*$\Delta$*SBS-nlsGFP* transgenes were integrated at the PBac{y+.attP}VK37 (22A3) site using phiC31-mediated integration. The *ato5'-AtoHalo* and *ato3'-AtoGFP* transgenes were integrated at the PBac{y+.attP}VK20 (99F8) and PBac{y+.attP}VK31 (62E1) sites. Injections were performed by BestGene Inc.

### S3. Immunostaining and smiFISH

Dissection of third instar larvae (in PBS 1x at room temperature), fixation (20 min in 4% paraformaldehyde in PBS 1x) and antibody staining in PBT (PBS 1X with Triton X-100 0.1%)

were performed using standard procedures. The following antibodies were used: goat anti-GFP (ab6673, Abcam), mouse anti-Hairy mAbs (clones 24/1, 28/2 and 33/5, from D. Ish-Horowicz), rabbit anti-Ato (from Y.-N. Jan and from D. Marenda), sheep anti-Ato (from A. Jarman), rat anti-Ecad mAb (DCAD2, DSHB) and guinea-pig anti-Sens (from H. Bellen). Secondary antibodies were from Jackson's laboratories.

Fluorescent in situ hybridization was performed as described previously (*35*). In brief, each smiFISH probe corresponds to duplexes between a set of gene-specific non-labelled primary oligonucleotides (probe set mix; equimolar mix of 23-29 different oligonucleotides with a 20 nt-long mRNA-binding moiety for a total length of 48 nt) which were annealed with a fluorescently labelled oligonucleotide (FLAP-X, 28 nt-long) coupled to Cy3, Alexa590 or Cy5. To obtain smiFISH probes, FLAP-X oligonucleotides were annealed with the probe set mix in Tris-HCl 50mM pH=7.5, NaCl 100mM, MgCl2 10mM using a thermocycler (85°C, 3 min; 65°C, 3min; 25°C 3 min). All oligonucleotides were obtained from IDT Inc. Dissected tissues were fixed 20 min in 4% paraformaldehyde in PBS 1x, washed in PBS 1x and permeabilized in PBS 1x Triton X-100 0.5%. Discs were then washed twice in SSC 2x with Urea 4M before being incubated overnight (ovn) with the smiFISH probe at 37°C in SSC 2x, Urea 4M, Dextrane 10%, Vanydyl complex 10mM, 0.15 mg/ml salmon sperm DNA. Following this hybridization step, discs were sequentially washed in SSC 2x Urea 4M (2x at 37°c), then in SSC 2x (2x at room temperature) and in PBT 1x. For double smiFISH/immunostainings, primary antibodies were added in the hybridization mix and incubated ovn at 37°c. Secondary antibodies were incubated in PBT following all washes.

Stained disks were mounted in 4% N-propyl-galate, 80% glycerol. Images were acquired using a confocal Zeiss LSM780 microscope with 40X (PL APO, NA DIC) and 63x (PL APO, N.A. 1.4 DIC M27) objectives.

### S4. *Ex vivo* culturing of eye discs

We used the explant culturing protocols described by (*37*) and (*38*) with minor modifications. Third instar larvae were briefly washed in 70% ethanol, rinsed in PBS 1x and dissected in dissection medium, i.e. Grace's medium (Sigma G9771) at pH 6.7 supplemented extemporaneously with 5% Fetal Bovine Serum (FBS), Penicillin/Streptomycin 0.5% (Sigma P4333) and 20nM 20-Hydroxyecdysone (Sigma H5142). Dissected eye imaginal discs still attached to mouth hooks and brain were transferred into a new dish with fresh medium for

further removal of the brain. They were then transferred via the mouth hooks to a magnetic imaging chamber (Chamlide CM-B25-1, LCI) with a drop of culture medium. Discs were then embedded using a fibrinogen-thrombin mix (Sigma 11424246 and 11407522), then incubated in dissection medium supplemented with Insulin (Sigma I9278 at 1.1 mg/ml).
For HaloTag labeling, 1.25µl of Janelia Fluor 646 HaloTag Ligand (200µM in DMSO; Promega GA1120) was added to the imaging medium. The Halo signal produced *in situ* by the proteins of interests was detected ~45 min after addition of the HaloTag ligand.

### S5. Live imaging

Movies were generally acquired by spinning disk microscopy using either a Leica DMRXA microscope equipped with a 40x (PL APO, N.A. 1.32 DIC M27) objective, a Yokogawa CSU-X1 spinning disk, a sCMOS Photometrics PRIM95B camera, 491/561/642 lasers and the Metamorph software, or a Nikon Ti2E microscope equipped with a 40X (N.A. 1.15 Water WD 0.6) objective, a Yokogawa CSU-W1 spinning disk, a sCMOS Photometrics PRIM95B camera and 488/561/640 lasers. Typically, we imaged the GFP/Halo and RFP signals every 5-10 mins and 1-5 mins, respectively, in a tissue volume with a typical depth of 40 $\mu$m ($\Delta z$=1.3$\mu$m). Under these conditions we were able to obtain reproducible dynamics of Ato expression during 6-7 hours of culture. A confocal Zeiss LSM780 microscope with 40X (PL APO, NA DIC) was occasionally used for live imaging (see Fig. S8C). Image acquisition was generally done at room temperature (~21-23°C).

### S6. Image analysis

#### S6.1. Ato kymographs

Kymographs of Ato expression were generated from maximum *z* projections of 4D image stacks, as follows. A rectangular region of interest (ROI) aligned with the MF was defined manually. For each time frame, the image was smoothed with a Gaussian filter with a standard deviation of 5µm, and a background level defined as the median intensity of the smoothed image within the ROI, then the spatial profile of Ato expression within the ROI was computed by projecting the signal above background onto the axis of the ROI.

S6.2. Period of Ato oscillations

The period of Ato oscillations was computed from the interval between successive peaks of Ato expression. For each analyzed movie, the spatial profile of Ato expression along the MF, as computed to generate kymographs, was averaged over a portion of the ROI (chosen to exhibit a strong signal and uniform timing of the Ato pulses) to generate a time-dependent signal. By linear interpolating the slope of the signal between successive movie frames, and taking the zero-crossings of the slope, the timing of the peaks was defined with sub-frame-interval resolution.

S6.3. 3D segmentation

3D segmentation of single nuclei labelled with nuclear RFP was performed using a custom-made program described in (*3*). The program generated a data frame with the spatial and temporal coordinates of the nuclei as well as the fluorescence intensities.

S6.4. Single cell tracking

After 3D segmentation we obtained a data frame with the position (x, y, z), time and Ato levels of the nuclei in the movie. Then we performed semi-automatic single cell tracking using the Fiji plugin Mastodon and tracks were assessed and manually corrected by visual inspection. Typically, the $\Delta t$ in the movies was 1 minute for non-sparse labelling and 3-6 minutes in the case of sparse labelling.

S6.5. Coarse-grained analysis of Ato dynamics

For the coarse-grained analysis of Ato dynamics, proneural clusters were first detected and tracked as follows. The maximum *z* projection of the Ato channel was filtered in space using a difference of Gaussians (taking the difference between the image filtered once with a Gaussian filter and twice with the same Gaussian filter) and smoothed in time using a Gaussian filter, to facilitate cluster detection and tracking in the presence of cell movement. Then, clusters were detected at each time step as local maxima of the filtered image (Fig. S2F). The standard deviations of the filters were chosen between 6 and 7.5 pixels in space, according to the spacing of the clusters, and between 10 and 15 minutes in time. Finally, the positions of the clusters in successive frames were connected into tracks using the Fiji plugin

TrackMate, which allowed for automatic tracking and manual correction and annotation when needed.

To analyze the distribution of Ato within proneural clusters and its time evolution, successive *z* stacks were segmented in 3D as described in (*3*) to detect nuclei and measure nuclear Ato-GFP levels. For each cluster and each time frame, each nucleus $i$ was assigned a weight $w_i$, defined as a Gaussian function of its distance to the cluster center in the *xy* plane, with a standard deviation of 7.5 pixels. The total Ato level in the cluster was then defined as the weighted sum of nuclear levels,

$$L = \sum_i w_i l_i \tag{S1}$$

where $l_i$ denotes the level in nucleus $i$, and the number of cells contributing to Ato expression within the cluster was defined in terms of the contributions $w_i l_i$ of each nucleus, as

$$N = \frac{(\sum_i w_i l_i)^2}{\sum_i (w_i l_i)^2} \tag{S2}$$

The above expression, which is exactly equal to the number of contributing cells when the contributions are binary ($w_i l_i = 0$ or 1), defines an effective cluster size when Ato levels are continuously distributed.

To compute averages of the above quantities within each row, as displayed in Fig. S2G,H, individual clusters were manually assigned to rows. To characterize the average dynamics of cluster resolution, based on pooling all the tracked clusters from two experiments, as shown in Fig. 1I, the clusters were registered in time according to the time when they reached the IG stage. This was defined from the time evolution of the total Ato level in the cluster (as defined above). The time course of Ato level in a cluster was filtered using a 90-min raised cosine window (one period of the function $[1 + \cos 2\pi t/90]/2$, chosen to approximate one period of the Ato signal), and the IG stage was defined as a peak of the filtered signal (with the manual assignment of clusters to rows serving to select the peak corresponding to the IG stage). By computing the filtered signal with a finer time step than the movie frame interval, the peaks were defined with sub-frame-interval resolution.

S6.6. Average Ato dynamics

An average time-dependent pattern of Ato expression was computed from successive images of proneural clusters that had been tracked and registered in time as described above. The images were rotated according to the orientation of the MF (from the manually defined rectangular ROI used for kymographs) then registered in space. To show where R8 cells emerge within IGs, we did not use the tracking of the clusters to align the images in space (by construction, this would show R8 cells emerging at the center of the clusters). Instead, particle image velocimetry (PIV) was used to track the 2D motion of the tissue from the average *z* projection of the nuclear signal and to map successive images of a cluster to a fixed reference frame, corresponding to the IG stage. When the time points at which the average pattern was computed mapped to non-integer numbers of frames in the original movies, the images included in the average were obtained by linear interpolation between consecutive movie frames.

S6.7. E(spl) kymographs

Kymographs of E(spl) expression were generated from maximum *z* projections of 4D image stacks, as follows. Movies were rotated to align the MF with the *y* axis, after which a ROI to be analyzed was defined manually as a range along that axis. For each time frame, the anterior boundary of the E(spl) expression domain was detected by smoothing the image with an anisotropic Gaussian filter, with a standard deviation of 4µm along the *x* axis and 16µm along the *y* axis, chosen to blur out the structure of the pattern along the MF, then fitting a smooth curve to the anterior edge of the region of high intensity, by minimizing a cost function that draws the curve towards regions of strong gradient while penalizing sharp variations in its profile. The spatial profile of E(spl) expression near the anterior edge of its expression domain was then obtained by smoothing the original image with a milder, anisotropic Gaussian filter, with a standard deviation of 5µm along the *x* axis and 2µm along the *y* axis, to blur our cellular-scale detail, and taking for each position along the MF the intensity of the smoothed image at a fixed distance (2.5µm) posterior to the detected domain boundary. For display, the kymographs were smoothed in time with a Gaussian filter with a standard deviation of one frame interval.

S6.8. Manual and automated annotation of E(spl) kymographs

Kymographs of E(spl) expression were manually annotated to label and connect successive peaks and troughs corresponding to a given A-P column, at an approximately stationary position along the MF. For each column, the local (high or low) phase of E(spl) expression was evaluated by taking the difference between the local E(spl) level (as measured in the kymograph) and the average level in adjacent columns. A time-dependent phase was then assigned to the column based on the zero-crossings of this relative E(spl) level, which identified alternations between teeth at staggered positions. The zero crossings were assigned discrete phase values, and the phase of intermediate time points was computed by linear interpolation. With one unit of phase chosen to correspond to the interval between successive rows of teeth, which matches the period of Ato pulses, the periodicity of E(spl) dynamics within columns is 2, and therefore the phase is defined modulo 2 (e.g. phases -1 and +1 correspond to the same stage). Zero-crossings with a negative slope, corresponding to the disappearance of a tooth (and emergence of teeth in adjacent columns), were assigned a phase -1/2 modulo 2, whereas zero-crossings with a positive slope, corresponding to the emergence of a tooth, were assigned a phase +1/2 modulo 2. With this choice, troughs in E(spl) expression, which coincide with the emergence of an IG, correspond to phase 0 modulo 2.

S6.9. Average E(spl) dynamics

An average time-dependent pattern of E(spl) was computed from successive images of manually labeled columns, registered in time according to their phase as defined above. Using PIV to correct for tissue motion, the images were mapped to a fixed reference frame, corresponding to the phase +1/2, and aligned according to the positions (at that stage) of the column along the MF and of the E(spl) domain boundary along the A-P axis. As with the average Ato dynamics, linear interpolation was used to generate images corresponding to non-integer numbers of frames. Note that in this procedure, a given image of a column in a given movie frame can appear in the average patterns for two different phases separated by 2 (the two images would be mapped to different reference frames and thus centered differently).

S6.10. Reconstruction of Ato/E(spl) dynamics from fixed samples

To reconstruct the dynamics of Ato and E(spl) from fixed samples, images of individual proneural clusters were ordered according to their stage in the progression of the pattern. To this end, average *z* projections of image stacks were manually annotated to label rows of IGs and emerging R8 cells. To include more positions and stages in the reconstruction, the manually labeled rows were complemented with staggered positions lying at the midpoints between successive positions in a row along the MF, and displaced by $\pm$5µm along the A/P axis, corresponding to the typical interval between rows. To assign a stage to each position, we leveraged the gradient in maturation along the MF, manually labeling discrete positions along the MF corresponding to peaks in Ato expression and E(spl) teeth (labeled as integer phases) and to the alternation between successive rows of teeth (labeled as half-integer phases), and defining the phase of intermediate positions by linear interpolation. For any given phase, an average pattern for that phase was then obtained by averaging the images of positions with a phase in a set interval (-.2 to +.2 phase units) around that phase.

S6.11. Analysis of Ato/E(spl) dynamics

For the analysis of movies of triply labeled eye disks, the E(spl) signal was processed and annotated as described above. To construct two-color kymographs showing the dynamics of Ato and E(spl), profiles of Ato expression were computed in the same way as E(spl) expression profiles, from the intensity of a smoothed image along a curve just posterior to the boundary of the E(spl) domain. To generate the kymograph in Fig. 3E, the Ato and E(spl) levels were detrended to correct for non-uniform signal intensity across the tissue. To this end, the kymograph was smoothed with a Gaussian filter with standard deviations of 10µm in space and 45min in time to measure the variations of the signal on scales larger than the relevant spatial and temporal scales of the pattern, after which signal levels in the kymograph were normalized to the smoothed levels. The average dynamics of Ato and E(spl) were reconstructed in the same way as for E(spl) alone, computing average patterns in the same way for the two channels.

S6.12. Kymographs of E(spl) and Ato/E(spl) within A-P columns

Kymographs showing the average dynamics of E(spl) and Ato/E(spl) within A-P columns were computed from images of individual columns registered in space and time as for the reconstruction of average expression patterns, by taking the average of the image levels along the direction of the MF, within a 2µm-wide strip running along the A-P axis (see Fig. S7).

S6.13. Analysis of single-cell E(spl) dynamics

Single-cell E(spl) time courses, from the tracking of sparsely labelled cells, were analyzed as follows. To correct for global variations in signal level over the duration of an experiment, E(spl) levels were normalized according to a time-dependent background level and signal amplitude, computed as the mean and standard deviation of the signal in the vicinity of the anterior boundary of the E(spl) expression domain (the signal included in this computation was obtained by multiplying the image levels by a Gaussian with a standard deviation of 75 pixels, centered on the boundary).

To register the single-cell tracks in time, A-P columns in each experiment were manually annotated and their time-dependent phase was computed as described above. Single-cell tracks were then registered according to local phase in the cycle of E(spl) dynamics, defined as the phase of the nearest A-P column. The phase of a column is defined modulo two, and the absolute phase of a track was chosen such that the initial onset of E(spl) expression in the cell (the time when its level first rose above background, shown as a dotted line in Fig. S16), corresponded to a phase in the interval between -2.5 and -.5. With this convention, the trough in E(spl) expression that follows the onset of E(spl) expression, which can be identified with the IG stage based on the relative dynamics of Ato and E(spl), corresponds to a phase around 0.

### S7. Tentative model for pattern propagation by negative templating

Our initial model of eye patterning by negative templating (Fig. S1, Movie S1) was formulated as a direct transposition of our earlier model for sensory organ patterning (*3*), with the same Notch-mediated cell-cell interactions but different initial and boundary conditions (as a deviation from (*3*), the models considered here do not incorporate a stochastic term in the dynamics). Specifically, the dynamics of cell $i$ is described in terms of a single state variable $u_i$ representing proneural activity, which varies according to the signal $s_i$ received by the cell:

$$\tau \frac{du_i}{dt} = f(u_i, s_i) - u_i \tag{S3}$$

with

$$f(u, s) = \sigma[2(u - s)] \tag{S4}$$

where $\sigma$ is the sigmoidal function

$$\sigma(x) = \frac{1 + \tanh 2x}{2} \tag{S5}$$

The signal $s_i$ received by the cell is

$$s_i(t) = s_0(x_i, t) + \sum_j c_{ij} D^*(u_j) \tag{S6}$$

In the above equation, the receding inhibitory front

$$s_0(x, t) = \sigma\left(\frac{x - x_0 - ct}{W}\right) \tag{S7}$$

with a width $W$ and velocity $c$ stands in for the traveling differentiation front that sweeps the eye disk. With a positive velocity $c$, the front travels in the direction of increasing $x$; in all figures and movies, this is oriented towards the left, to match the orientation of the A-P axis in experimental images.

As in (*3*), the coupling between cells $i$ and $j$ is a function of their distance

$$c_{ij} = e^{-\frac{d_{ij}^2}{2l^2}} \tag{S8}$$

and the active ligand level in a cell depends on its cell state through

$$D^*(u) = \left(a_0 + \frac{3u^3}{2 + u^2} a_1\right) u \tag{S9}$$

As in (*3*), simulations were run on a fixed lattice of cells, with disordered cell arrangements generated using a vertex model of epithelial tissue dynamics.

The simulation shown in Fig. S1 and movie S1 was run on an array of 1024 cells in a 32x32 box (corresponding to unit density) with periodic boundary conditions along the $y$ axis (parallel to the front). The simulation was initialized with a template comprised of 4 R8 cells, with fixed $u = 1$, representing one row in the pattern. To mimic inhibitory signaling from R8 cells in earlier rows and prevent proneural activation in cells posterior to the template, an additional source of inhibition at $x = 0$ and with a range $l$ was added to the extrinsic field $s_0$:

$$s_0(x, t) = e^{-\frac{x^2}{2l^2}} + \sigma\left(\frac{x - x_0 - ct}{W}\right) \tag{S10}$$

The template R8 cells were chosen as the closest cells to evenly spaced points, at a position $x = \Delta x$ along the direction of front motion and separated by $\Delta y$ along the $y$ axis, with $\Delta x$ chosen to approximately match the spacing of R8 rows in simulations. The initial position $x_0 = 2\Delta x$ of the front was chosen such that proneural activation anterior to the template began shortly after $t = 0$ (see Table S1 for parameter values).

### S8. Main model

#### S8.1. Signal-dependent differentiation

Our main model elaborates on our earlier model (*3*) by incorporating an additional cell state variable $m$ representing the maturation of a cell from undifferentiated to differentiated and allowing the dynamics of the cell to vary as it matures:

$$\frac{dm_i}{dt} = f_m(m_i, s_i) \tag{S11}$$

$$\tau \frac{du_i}{dt} = f(m_i, u_i, s_i) - u_i \tag{S12}$$

The form of the function $f_m$ is chosen such that signaling is required to initiate differentiation, after which $m$ increases at an approximately constant, unit rate, effectively measuring time since differentiation was triggered. To this end, we take

$$f_m(m, s) = -\frac{\partial}{\partial m} V(m, s) \tag{S13}$$

where the potential

$$V(m, s) = -m + (m_c \cosh^2 1) h(s) \sigma\left(\frac{m}{m_c} - \frac{1}{2}\right) \tag{S14}$$

is comprised of a signal-dependent sigmoidal step overlaid on a uniform slope, yielding

$$\frac{dm_i}{dt} = 1 - h(s_i) \frac{\cosh^2 1}{\cosh^2\left(2\frac{m_i}{m_c} - 1\right)} \tag{S15}$$

The term

$$h(s) = \sigma(1 - k_s s) \tag{S16}$$

controls the size of the step, which decreases with the signal level (see Fig. S11D). Cells are initialized with $m = 0$. In the absence of signaling, $h \approx 1$ and cells remain in the minimum of the potential near $m = 0$. For sufficiently large signal levels, such that $h < h_c = 1/\cosh^2 1 \approx 0.4$, the minimum disappears and $m$ increases, at a rate that becomes independent of the signal and tends to one for sufficiently large $m$. The parameterization is chosen such that $m = 0$ and $m = m_c$ are fixed points when $h = 1$, such that a cell with $m > m_c$ is committed to differentiating.

### S8.2. Maturity-dependent dynamics

The modulation of proneural activity and its response to signaling as cells mature is encoded in the function $f(m, u, s)$, cf. eq. S12. In its simplest form, this incorporates the transition between an early regulation of proneural activity, which is promoted by Notch signaling, and a late regulation, which depends on the balance between self-activation and inhibition by Notch, as in (*3*) (the incorporation of commitment is described in a separate section). To this end, we introduce the functions

$$\chi_{\text{early}}(m) = \sigma\left(\frac{m_{\text{early}} - m}{\tau_{\text{early}}}\right) \tag{S17}$$

$$\chi_{\text{late}}(m) = \sigma\left(\frac{m - m_{\text{late}}}{\tau_{\text{late}}}\right) \tag{S18}$$

and write

$$f(m, u, s) = a_{\text{early}}\chi_{\text{early}}(m) f_{\text{early}}(s) + \chi_{\text{late}}(m) f_{\text{late}}(u, s) \tag{S19}$$

where $a_{\text{early}}$ parameterizes the level of proneural activity in the early regime, and the functions $f_{\text{early}}(s) = \sigma(4s - 2)$ and $f_{\text{late}}(u, s) = \sigma(2(u - s))$ represent the early and late regulation of proneural activity. The steady states of proneural activity as a function of signal level in the two regimes are plotted in Fig. S11E, showing a monotonous increase in the early regime, whereas the late regime exhibits bistability in an intermediate range of signal levels, as required for the progressive restriction of proneural clusters through lateral inhibition (*3*).

### S8.3. Maturity-dependent signaling

Our model depends on a transient increase in the signaling range from cells in resolving IGs. The gradual extension of E(spl) teeth observed in experiments is modeled through a maturity-dependent length

$$l_t(m) = l_{\min} + \sigma\left(\frac{m - m_{l_{\min}}}{m_{l_{\max}} - m_{l_{\min}}}\right)(l_{\max} - l_{\min}) \tag{S20}$$

while their subsequent disappearance is represented by a maturity-dependent amplitude

$$a_t(m) = \sigma\left(\frac{m_t - m}{\tau_t}\right) \tag{S21}$$

The weight of the signal reaching cell $i$ from a tooth with its source in cell $j$ is described by the coupling coefficient

$$c_{ij,t}(m_j) = e^{-\frac{(y_i-y_j)^2}{2w_t^2}} \sigma\left(\frac{x_j - x_i + X}{\Delta X}\right) \sigma\left(\frac{x_i - x_j - l_t(m_j)}{\Delta X}\right) \quad \text{(S22)}$$

In the above expression, $w_t$ is the width of teeth along the MF and $W_t$ the width of the step in signal level along the A-P axis at the ends of the tooth. The term in $X$, which delimits the tooth in the posterior, essentially plays no role in the dynamics; together with $\Delta X$, it is chosen to yield a profile similar to the basal signaling range from Eq. S8.

Finally, the transition from transient signaling in teeth to persistent signaling is described by

$$s_i = \sum_j \max\{c_{ij,t}(m_j)a_t(m_j), c_{ij}\}D^*(m_j, u_j) \quad \text{(S23)}$$

where

$$D^*(m,u) = \sigma\left(\frac{m - m_{D^*}}{\tau_{D^*}}\right) D^*(u) \quad \text{(S24)}$$

In these expressions, the coupling coefficients $c_{ij}$ and the function $D^*(u)$ from eqs. S8 and S9 represent the persistent signaling from R8 cells, whereas $D^*(m, u)$ incorporates a dependence of signal production on cell maturity that restricts mutual inhibition to IGs.

#### S8.4. Commitment

A finite time window for fate specification is built into the model by letting the dynamics become independent of the signal in more mature cells. To do so, the signal $s$ in the regulation function $f_{\text{late}}(u, s)$ in Eq. S19 is replaced by an effective inhibitory input $h(m, s)$

$$f_{\text{late}}(u, s) = \sigma[2(u - h(m, s))] \quad \text{(S25)}$$

where $h(m, s)$ tends for large $m$ to a signal-independent value of $1/2$, yielding bistable dynamics.

Based on the phenotype of a temperature-sensitive Notch mutant, when Notch function is suppressed, Ato expression persists in IGs but is not restored in earlier rows (*3*), suggesting that non-R8 cells are committed shortly after they are excluded from the proneural group. On the other hand, the gradual resolution of IGs (Fig. 1I) suggests that the remaining proneural cells should remain responsive to inhibition. To incorporate commitment into the model while allowing for such differences in responsiveness, the response to high and low signal levels is modulated separately in time. To this end, maturity-dependent sensitivities $\xi_{\text{low}}(m)$ and $\xi_{\text{high}}(m)$ are introduced, and we set

$$h(m,s) = \frac{1-\xi_{\text{low}}(m)}{2} + \int_0^s ds' \left[\xi_{\text{low}}(m)\sigma(1-2s') + \xi_{\text{high}}(m)\sigma(2s'-1)\right] \quad \text{(S26)}$$

This expression is chosen such that the slope of $h(m,s)$ crosses over from $\xi_{\text{low}}(m)$ to $\xi_{\text{high}}(m)$ around $s = 1/2$, with a value around 1/2 at the crossover. As shown in Fig. S11F,G, when the sensitivity $\xi_{\text{low}}$ alone is reduced, cells with low proneural activity are committed while cells with high proneural activity remain responsive to inhibition. Eventually, as $\xi_{\text{high}}$ also tends to 0, the dynamics becomes signal-independent.

The two functions $\xi_{\text{low}}(m)$ to $\xi_{\text{high}}(m)$ describing these successive transitions are sigmoidal step functions

$$\xi_{\text{low}}(m) = \sigma\left(\frac{m_{\text{low}} - m}{\tau_{\text{low}}}\right) \quad \text{(S27)}$$

and

$$\xi_{\text{high}}(m) = \sigma\left(\frac{m_{\text{high}} - m}{\tau_{\text{high}}}\right) \quad \text{(S28)}$$

With commitment incorporated in this way, signal production can be phased out in the more mature cells. Using the same parametrization for this as for the second of the above transitions, after which all cells are irresponsive to signaling, we take

$$D^*(m,u) = \sigma\left(\frac{m - m_{D^*}^{\text{ON}}}{\tau_{D^*}}\right)\sigma\left(\frac{m_{\text{high}} - m}{\tau_{\text{high}}}\right)D^*(u) \quad \text{(S29)}$$

#### S8.5. Template

Since pattern propagation in the model depends on signaling from row *n* to row *n*+2, the model is initialized with a template comprised of two rows, identified in the following as rows -1 and 0 (Fig. S11H). The template is specified by designating cells to represent the R8 cells in those two rows. These are chosen as the closest cells to regularly spaced points with coordinates of the form $\{x_0 - \Delta x, (k + 3/4)\Delta y\}$ for row -1 and $\{x_0, (k + 1/4)\Delta y\}$ for row 0, with $k$ integer. In these expressions, $x_0$ is the position of row 0 along the $x$ axis, and $\Delta x$ and $\Delta y$ are the spacing between and within rows, respectively. To define the signal they produce, the template R8 cells in row $r$ are assigned a fixed state of maximal proneural activity, $u = 1$, and a maturity $m(t) = t - r$, representing the maturity of cells that initiated differentiation around $t = r$ (the timing is chosen such that the first row following the template, row 1, engages in differentiation around $t = 1$, making $t = 0$ an adequate starting time for simulations).

The onset of differentiation in the rows that follow the template, rows 1 and 2, depends on the activating signal from rows -1 and 0, but also on which cells have already initiated differentiation as part of rows -1 and 0 and are therefore excluded from being part of rows 1 and 2. To define this, a preliminary simulation of the equation for cell maturity (Eq. S11) is run with signal sources at a distance $2\Delta x$ posterior to each R8 cell in the template, representing R8 cells in earlier rows,-3 and -2. The signal produced by each source is computed as the signal of a cell with full proneural activity ($u = 1$) and with a maturity $m(t) = t + 3$ for sources representing signaling from row -3, and $m(t) = t + 2$ for sources representing signaling from row -2. This preliminary simulation is run for a sufficient time to include all significant signaling from the sources, after which the cells that have crossed the threshold $m = m_c$ are deemed to have engaged in differentiation. In the full simulation, these cells, with the exception of the template R8 cells, are kept in a fixed state with no proneural activity ($u = 0$), representing committed non-R8 cells.

#### S8.6. Uniform template

To initialize pattern propagation with a uniform template, differentiation is triggered in a narrow strip of cells in the interval $x_0 < x < x_0 + \Delta x$, with a gradient in initial maturity to bias the selection of the first R8 cells to the posterior of the strip. Specifically, cells with $x < x_0$ are excluded from patterning (with fixed $u = 0$), whereas cells with $x > x_0$ are initialized with $m = 1 - x/\Delta x$ if $x < x_0 + \Delta x$ and $m = 0$ if $x > x_0 + \Delta x$. This initial condition, which can be likened to the state of cells within IGs, but rendered uniform along the MF, results in relatively uniform activation in a file of cells before mutual inhibition sets in (Fig. S11I).

#### S8.7. Parameter values

For those parameters that already appeared in our earlier model of lateral inhibition patterning (*3*), the same values were kept here; as an exception, the time scale $\tau$ was set to a smaller value, since one unit of time was chosen here to correspond to the period of Ato oscillations ~ 90min, when it was chosen to correspond to approximately one hour in (*3*). Elaborating on our earlier model to incorporate maturity-dependent dynamics and signaling required that we introduce additional parameters. As detailed below, the values of many of these new parameters are constrained by the observed dynamics of Notch signaling and proneural activity. Elsewhere, we found that default values (e.g. time scales commensurate with the period of the dynamics) were adequate (see Table S2 for the full list of parameter values). Based on our observation of E(spl) along A-P columns (see the kymographs in Fig. S7D,E), the anterior boundary of Notch activity progresses discontinuously, advancing from the IG in row *n* to the IC in row *n*+2 (or twice the spacing between rows) in about one unit of phase (corresponding also to one period of Ato), before Notch activity subsides for the next unit of phase. The model parameters controlling time-dependent signaling production were chosen to mimic this progression. Likewise, the parameters controlling the transition between early and late regulation were chosen to reproduce the timing of Ato expression in ICs and IGs. Our numerical experiments indicated that the signaling range $l$ should be slightly larger than the width $w_t$ of the teeth for stable pattern propagation (thus $w_t$ was chosen to be slightly less than the value of $l$ taken from (*3*)); as seen in simulations with a reduced signaling range (cf. Fig. S13B), multiple R8 cells could otherwise emerge from a single IG. As discussed above (see Commitment), the timing of commitment to a non-R8 fate (controlled by $m_{\mathrm{low}}$) was chosen to occur shortly after cells are excluded from the proneural group. In principle, the model allows for different time scales for the different transitions in the dynamics as cells mature. However, for simplicity, we set a single value for all those parameters; we reasoned that this value should be smaller, but not much smaller, than the period of the dynamics. As an exception, we set a longer time scale for the phasing out of signaling (controlled by $\tau_{\mathrm{high}}$), since our observations did not show a sharp down-regulation of Notch activity following R8 selection (as compared with the down-regulation of Notch activity in the interval between IC and IG, cf. Fig. S7D,E). As regards the dynamics of the maturity variable $m$, the sensitivity $k_s$ was chosen to be sufficiently large that weaker signaling in the teeth tips is sufficient to initiate differentiation, and the value of $m_c$ sufficiently small that the time after which cells are irreversibly engaged in differentiation is small (since this should occur in a fraction of one period of the dynamics).

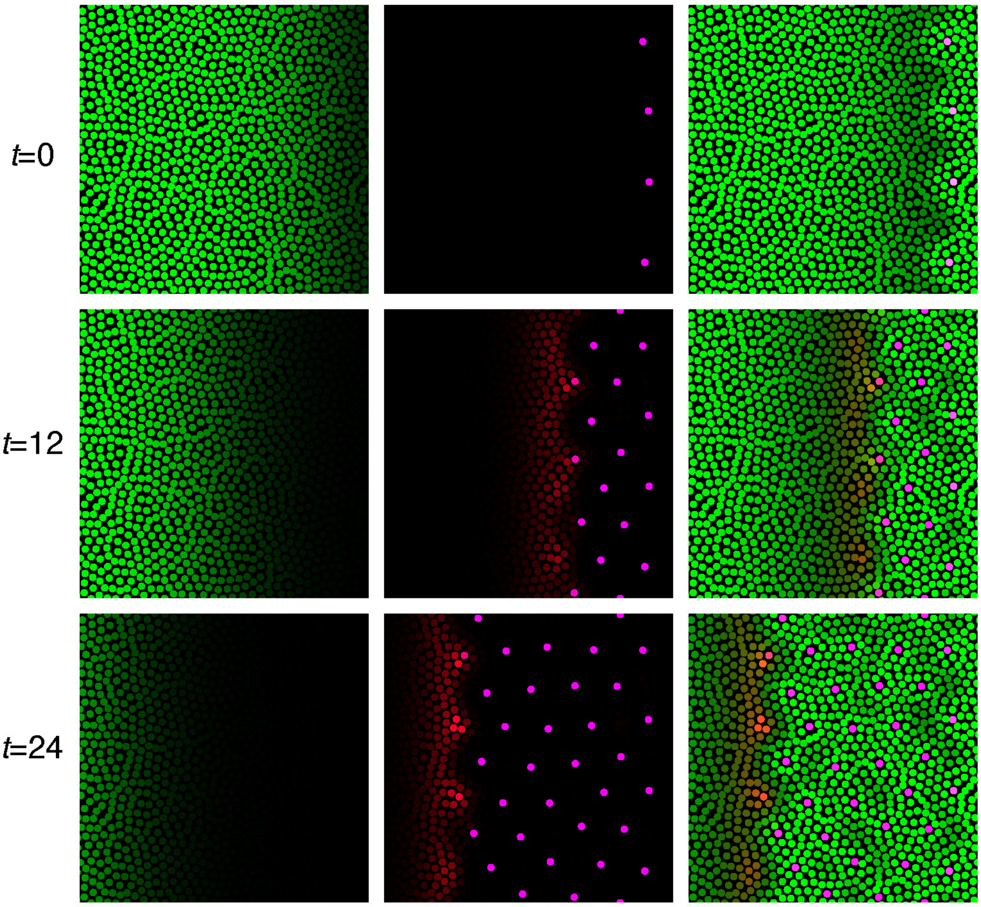

**Figure S1: A tentative model for pattern propagation by negative templating**
Snapshots from a simulation of a tentative model for eye patterning by negative templating, which combines cell-intrinsic bi-stability and short-ranged inhibition as in (*3*) with a receding inhibitory front (green, inhibitory signal from the front and cell-cell signaling; red/magenta, intermediate/high values of the cell state variable $u$). The left-hand column shows the inhibitory front alone, while the middle column shows the cell states alone (initialized with a regular template at $t = 0$), and the right-hand column the cell states along with the total inhibitory input to the cell dynamics, integrating the inhibitory front and cell-cell signaling.

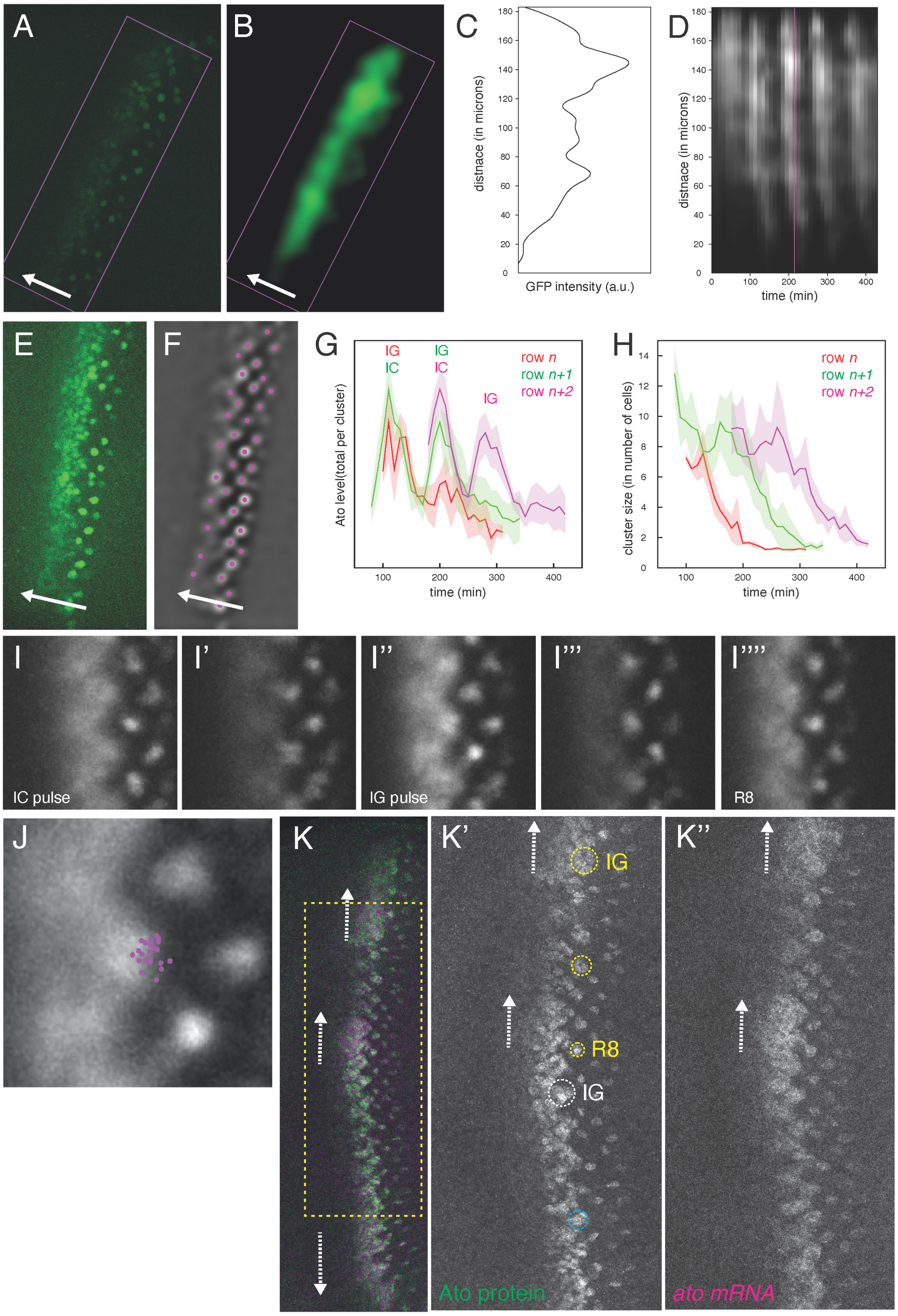

**Figure S2: Analysis of Ato dynamics**

(**A**-**D**) Ato kymographs. A rectangular ROI captured the Ato signal at the MF (A, maximal projection; B, Gaussian filtered; AP axis indicated by an arrow). The smoothed Ato signal projected along the ROI long axis (C) was used to generate kymographs (D; time point shown in A-C indicated in magenta). See also Methods.

(**E**,**F**) The Ato signal (maximal projection, E) was smoothed in space and time and local maximum intensity peaks were identified (F), tracked over time and registered in time according to when they reach the IG stage.
(**G**,**H**) Total level of Ato in nuclei contributing to the clusters in three consecutive rows (G; mean and standard deviation values shown for each row) and number of nuclei contributing to the clusters (cluster size, H). See also Methods.
(**I**-**J**) Average dynamics of Ato over 3 pulses (I, IC stage; I'', IG stage; I'''', R8 stage; n=22 clusters from 2 movies). The R8 positions (dots) were mapped back to the IG stage using PIV, showing that R8s are selected from the posterior-most IG cells (J).
(**K**-**K''**) Pulses of endogenous Ato protein (green) and *ato* mRNA (magenta) accumulation at the MF. The white arrows show the direction of the TWs (based on the spatial progression of patterning from IG to R8 within a row (yellow, K'; panels K',K'' show the region boxed in K). Between pulses, lower levels of Ato were detected in ICs (arrowheads) and resolving IGs. The strict overlap between mRNA and protein levels indicated that both Ato proteins and *ato* mRNAs are short-lived relative to the period of the Ato oscillations.

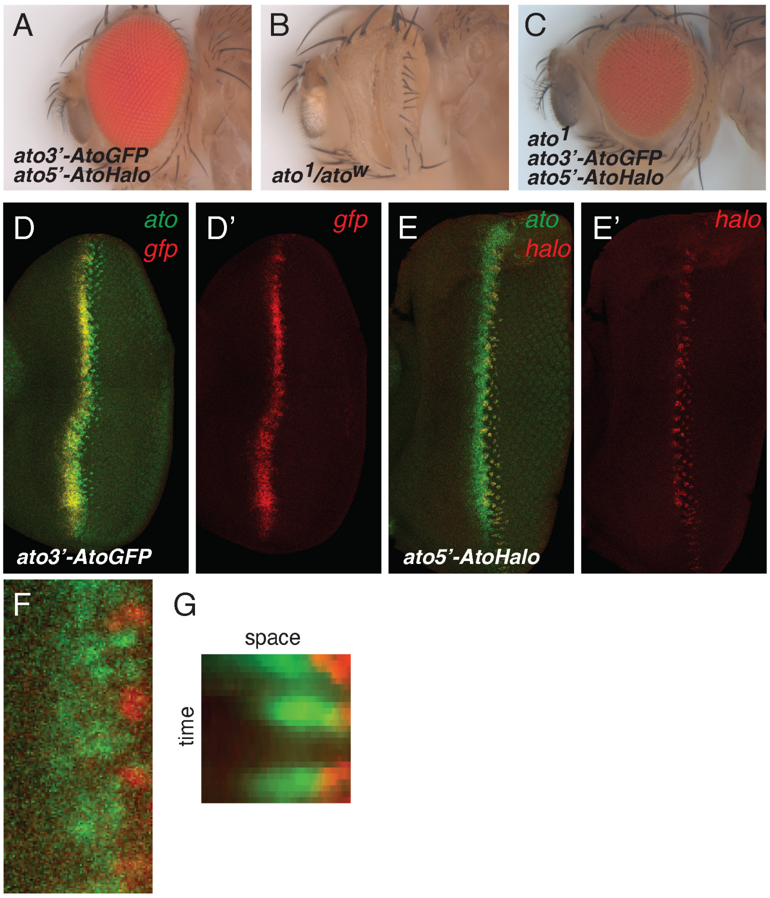

**Figure S3: Expression and function of ato3'-AtoGFP and ato5'-AtoHalo**

(**A**-**C**) The co-expression of AtoGFP and AtoHalo under the control of the *ato3'* and *ato5'* enhancers, respectively, was sufficient to rescue the *ato*$^{1}$ mutant adult eye phenotype (C; compare with wild-type (A) and *ato* mutant (B) control flies).

(**D**-**E'**) The *ato3'* enhancer (*gfp* mRNA, red in D,D') is active in anterior MF *ato*-expressing cells (*ato* mRNA, green) whereas the *ato5'* enhancer (*halo* mRNA, red in E,E') is active in IGs and R8s.

(**F**,**G**) Expression of AtoGFP (ato3'-AtoGFP, green) and AtoHalo (ato5'-AtoHalo, red) at t=220 (see Fig. 2C) in the MF region used for kymograph analysis (F). The GFP and Halo signals were projected along the A-P axis to produce a kymograph (G) and a plot (Fig. 2D).

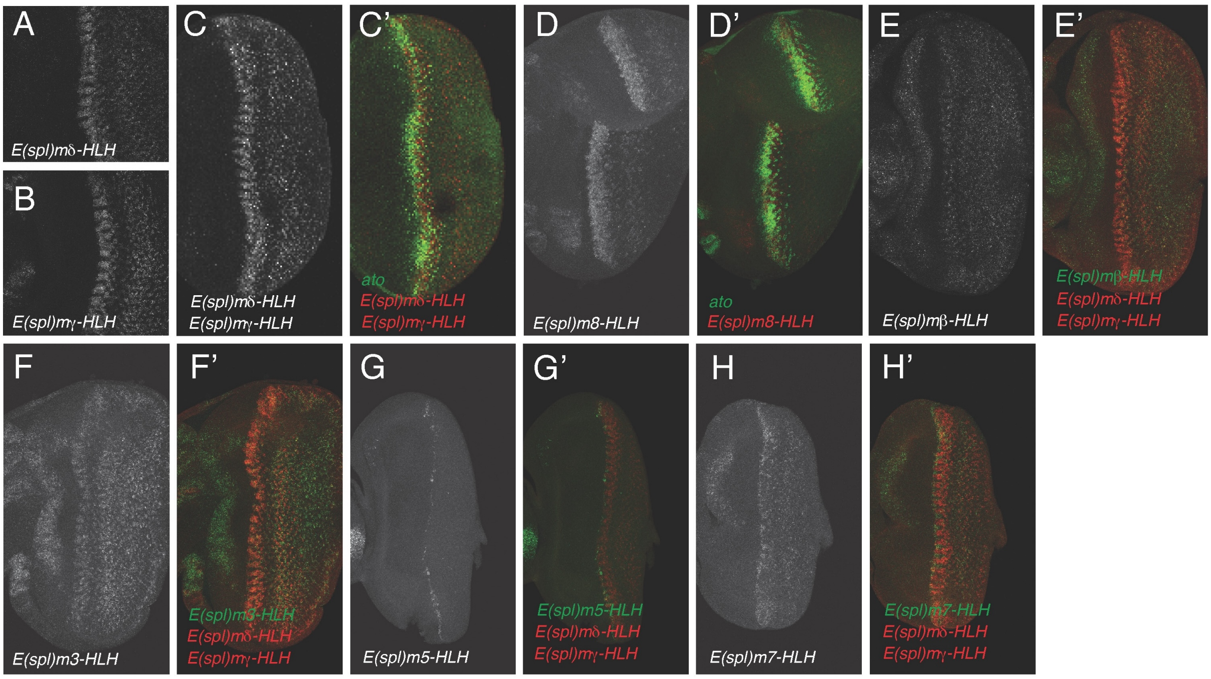

**Figure S4: Expression patterns of *E(spl)-HLH* genes**

(**A**-**H'**) mRNA accumulation of the *E(spl)-HLH* genes relative to *ato* (C',D') or to $m\delta$ and $m\gamma$, noted *E(spl)* here (C, E'-H')

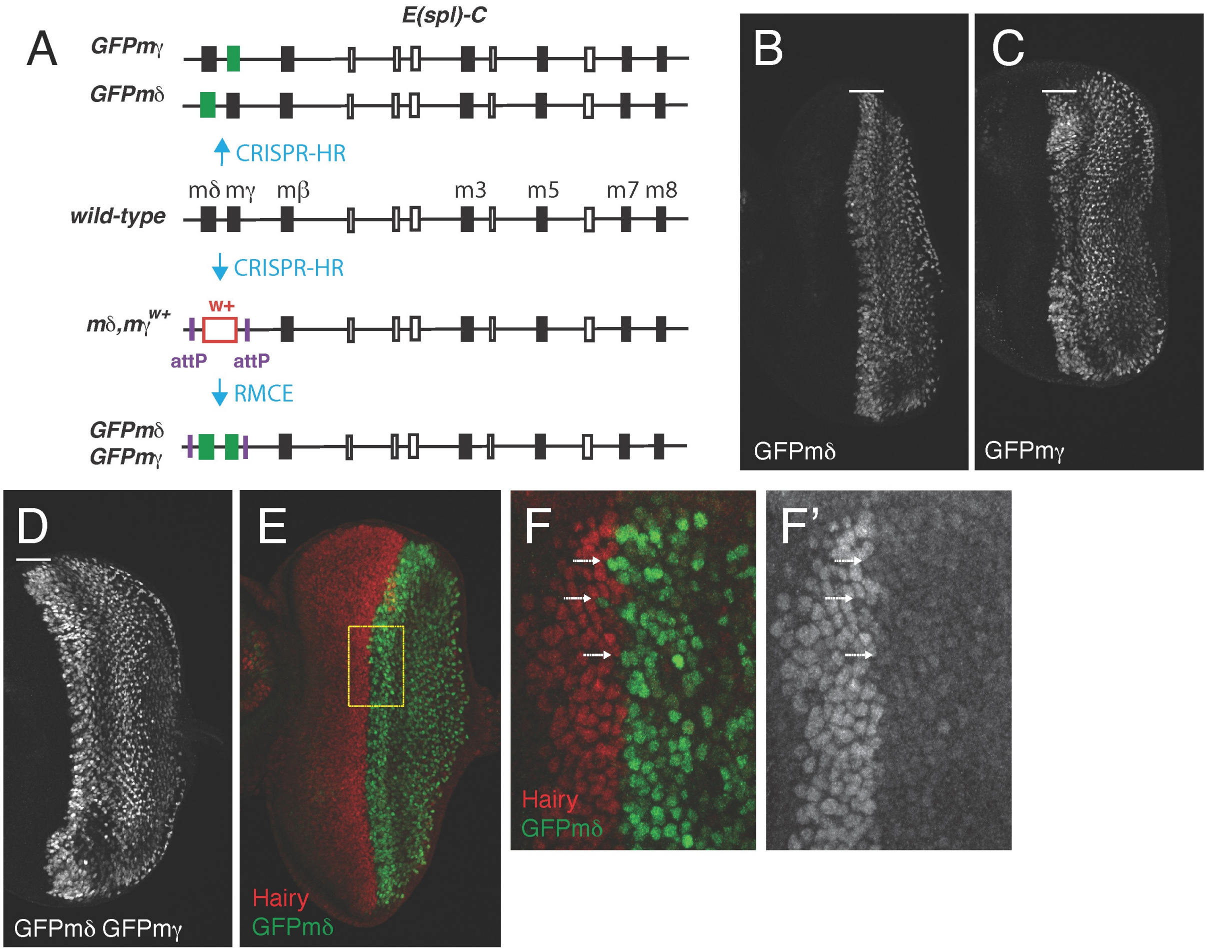

**Figure S5: GFP tagging of *E(spl)-HLH* genes**

(**A**) Genome engineering of the *E(spl) Complex* locus. CRISPR-HR was used to GFP-tag individually mδ and mγ, as well as to replace a genomic fragment encoding these two genes by a *white* (*w+*) marker flanked by attP sites. RMCE was then used to replace this marker by a modified version of this fragment encoding mδ and mγ both GFP-tagged.

(**B**-**D**) Anti-GFP staining of GFP-tagged mγ, mδ and mδ mγ eye discs.

(**E**-**F’**) Hairy (red) accumulates anterior to E(spl) proteins (GFPmδ, green) in strictly complementary manner. Note that the E(spl) teeth coincide with indentations along the Hairy expression boundary (F,F’, high magnification views of the region boxed in G).

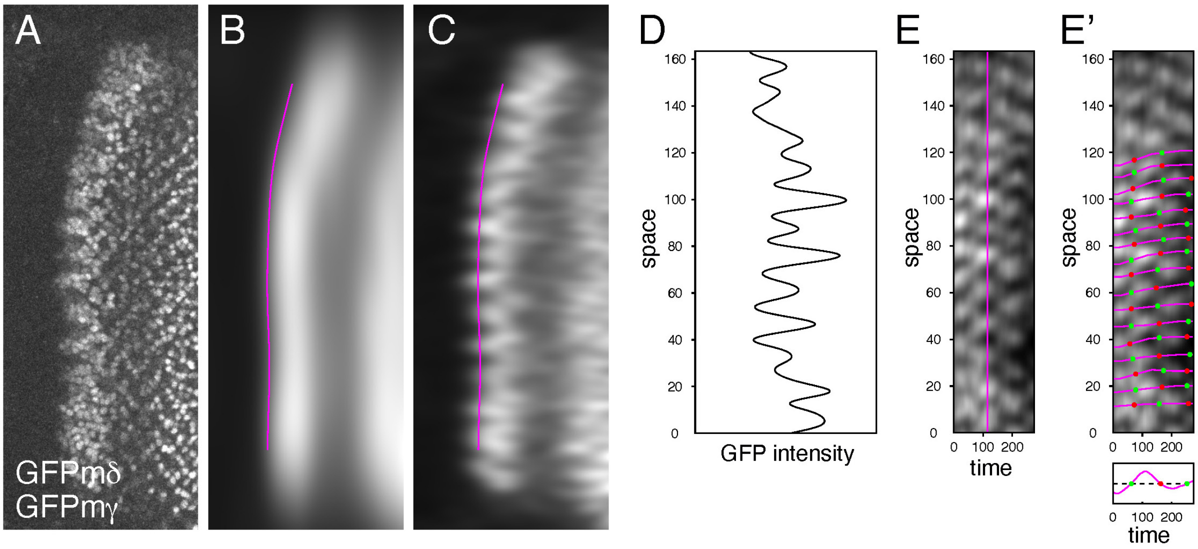

**Figure S6: Kymograph analysis of E(spl)**

(**A**-**E**) E(spl) kymographs. Starting with a maximum projection of the GFP signal (A), the anterior boundary of E(spl) expression, blurred using an anisotropic Gaussian filter, was automatically detected (magenta line, B). The spatial profile of E(spl) expression close to this line was then obtained by smoothing the signal with a milder anisotropic Gaussian filter (C) and computing the intensity along the line (D) to produce a kymograph (E; the time point shown in A-D is indicated in magenta). See Methods for details.

(**F**) Successive peaks and troughs of E(spl) were connected within each column (top panel, magenta lines) and the local E(spl) level was measured along this line. Discrete 'phase' values were defined at zero-crossing points (bottom panel). See Methods for details.

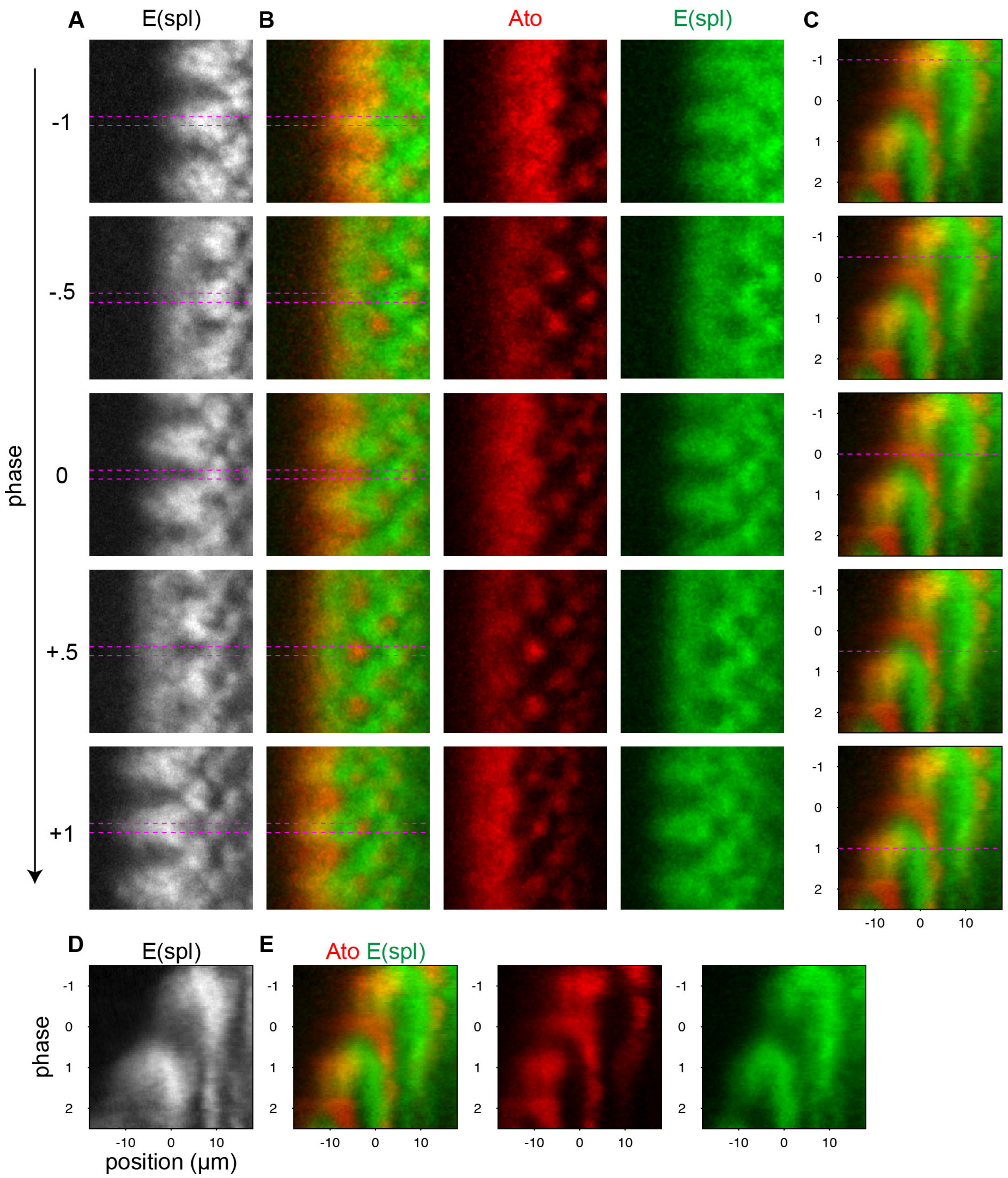

**Figure S7: Average dynamics of Ato and E(spl)**

(**A**) Average dynamics of GFPmδ GFPmγ, from n=15 positions in one movie, registered in space using PIV and in time according to the local phase of the dynamics (see Methods). Dashed lines indicate the region used to generate the kymograph in D.

(**B**-**C**) Average dynamics of HaloAto and GFPmδ, from n=16 positions in one movie, registered in space using PIV and in time according to the local phase of GFPmδ. Dashed lines in B indicate the region of the average pattern used to generate the kymograph in E and reproduced in C. For each of the shown phases, the kymograph is reproduced in C with a line indicating the phase.

**(D-E)** Kymographs summarizing the dynamics of E(spl) (D; GFPm$\delta$, GFPm$\gamma$), and of Ato and E(spl) (E) within A-P columns.

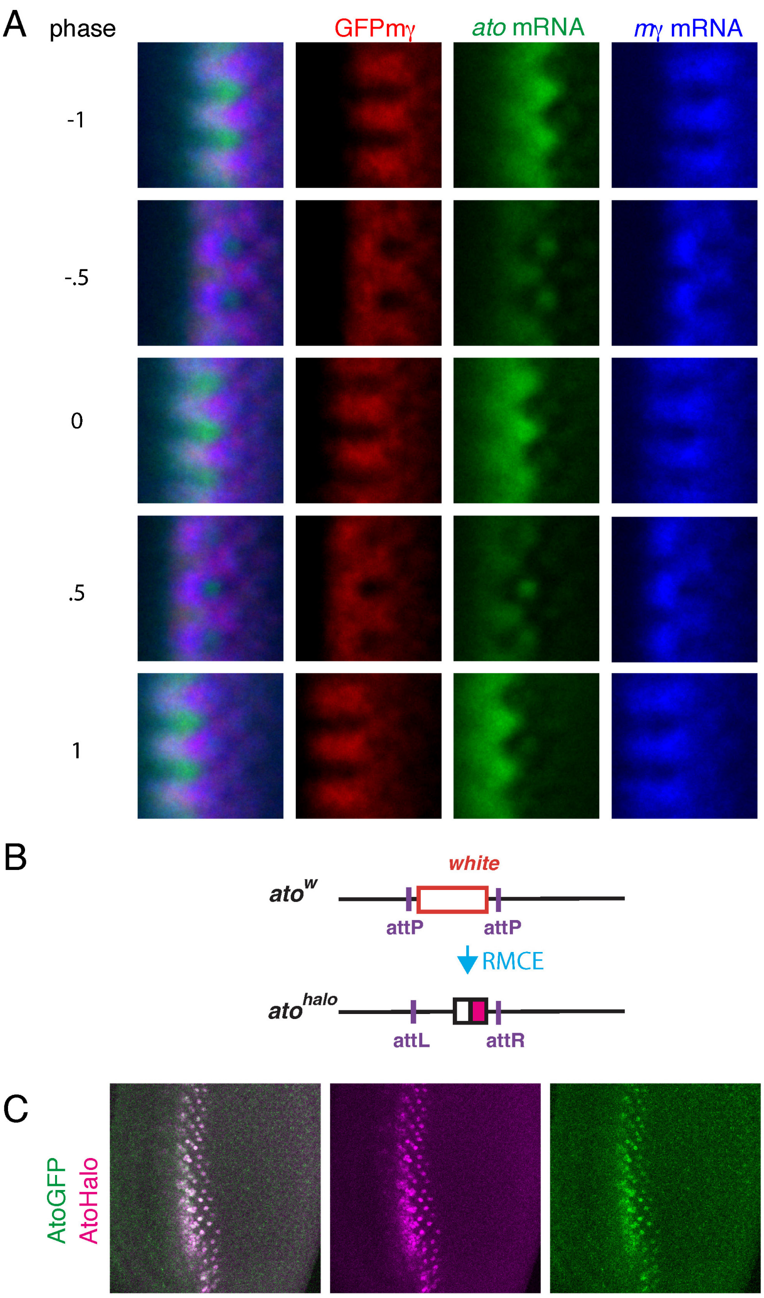

**Figure S8: Relative dynamics of E(spl) and Ato**

(**A**) The average pattern of *ato* mRNA, *GFPm*$\gamma$ mRNA, and GFPmγ protein at successive phases in the dynamics was reconstructed from n=96 positions in 4 fixed eye discs, registered in space and time based on the manual annotation of proneural cluster positions and of the phase of the dynamics along the MF (see Methods).

(**B**) RMCE-mediated engineering of the *ato* locus to produce the *ato*$^{halo}$ allele that encodes AtoHalo.

(**C**) Snapshot from a movie showing that AtoHalo and AtoGFP have similar expression dynamics. This in turn suggests that the labeling kinetics of AtoHalo and the maturation kinetics of AtoGFP are similar, and that both proteins are similarly short-lived.

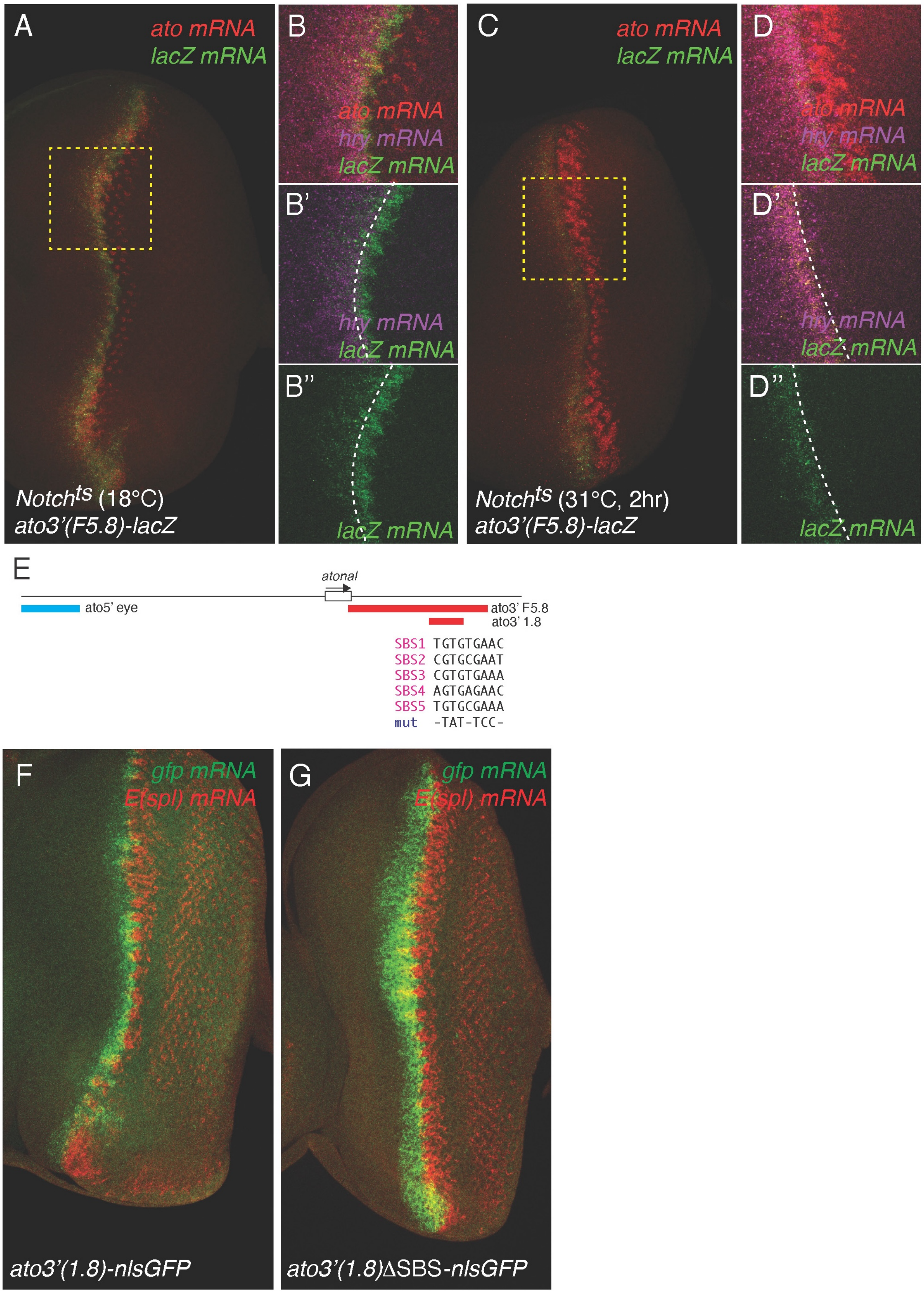

**Figure S9: Notch regulates the *ato3'* enhancer via Su(H)**

(**A**-D**''**) Conditional loss of *Notch* activity ($N^{ts}$ male larvae, 2h at 31°C; C-D'') resulted in elevated *ato* mRNA accumulation in two rows of IGs (*ato*, red) which failed to resolve due to disrupted Notch signaling relative to controls (control $N^{ts}$ male larvae, raised at 18°C; A-B''; B-B'' and D-DB'' are high magnification views of the boxed area in A and C, respectively). The activity of the *ato3'* enhancer (*ato3'(F5.8)-lacZ*, green) was strongly down-regulated in the Hairy (Hry)-negative cells of the MF (*hry* mRNAs, magenta), indicating that Notch signaling is required to up-regulate *ato3'* enhancer activity. In contrast, the activity of the *ato3'* enhancer in the *hry*-positive pre-proneural domain did not appear modified upon loss of *Notch* function (the limits of the *hry* expression domain are indicated by white dotted lines).
(**E**) representation of the *ato* locus showing the positions of the *ato5'* (*16*) and *ato3'(1.8)* enhancers. Five putative SBS were identified in the *ato3'*(*1.8)* enhancer (*39*), together with the mutations introduced to disrupt these sites.
(**F,G**) Mutation of the *ato3'(1.8*)∆*SBS* resulted in increased *ato3'(1.8)* enhancer activity (*gfp* mRNA, green; compare G with the control in F), indicating that Su(H) acts, at least in part, as a transcriptional repressor in this context.

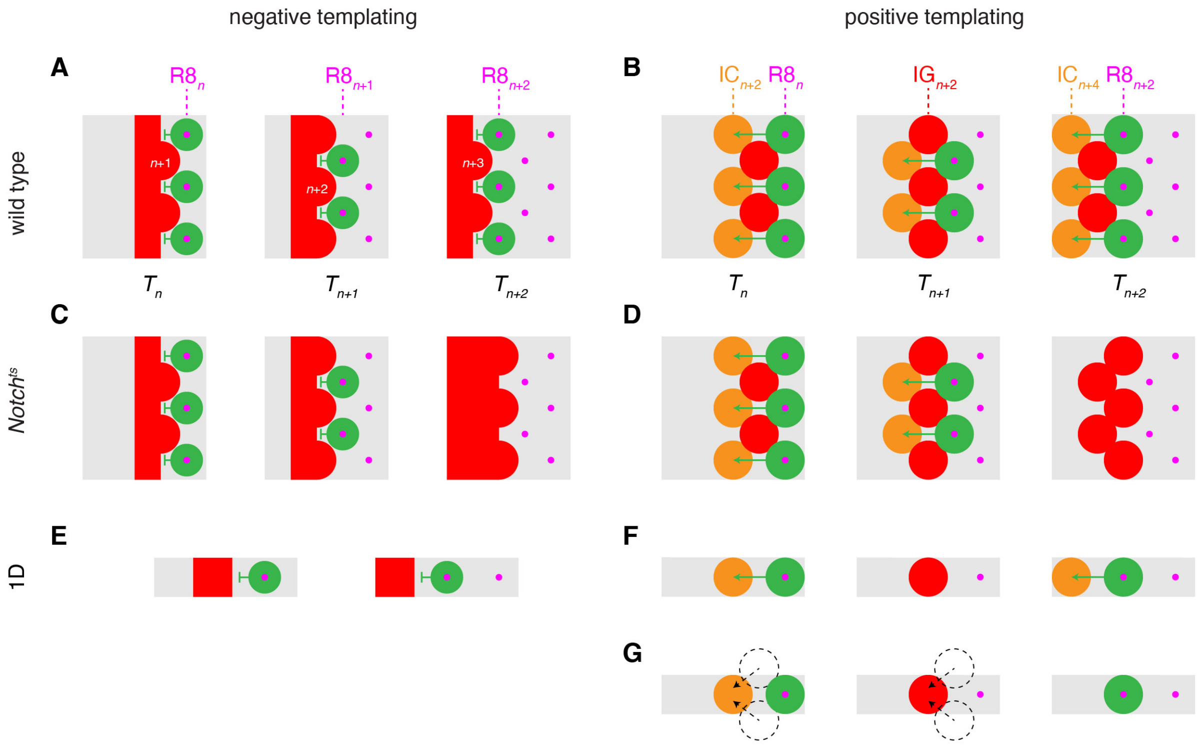

**Figure S10: Alternative models for pattern propagation and predicted outcomes in different conditions.**

(**A**) In a negative templating model, each row of R8s (e.g. row *n*) provides a template for the next row of IGs (*n*+1).

(**B**) In the positive templating model we propose, spatial information is relayed from emerging R8s in row *n* to ICs in row *n*+2.

(**C**) In a negative templating model, spatial structure along the MF should be lost upon conditional Notch inactivation (here, between times $T_{n+1}$ and $T_{n+2}$), except at its posterior edge where cells are committed to a non-R8 fate.

(**D**) In our positive templating model, spatial information is retained upon conditional Notch inactivation, with Ato expression being restricted to cells in which earlier signaling has initiated differentiation.

(**E**,**F**) Pattern propagation in 1D. Negative and positive templating models can both produce 1D spacing patterns, but predict different dynamics: in a negative templating model (E), Ato expression in clusters of cells and R8 emerge recur with the same temporal frequency, whereas our positive templating model predicts that non-R8-forming ICs should alternate with R8-forming IGs (F).

(**G**) A model in which the initiation of differentiation in ICs in row *n*+2 depends on signaling from row *n*+1 (arrows) is ruled out by the experimentally observed 1D propagation (in which row *n*+1 is absent, as symbolized by the dashed circles).

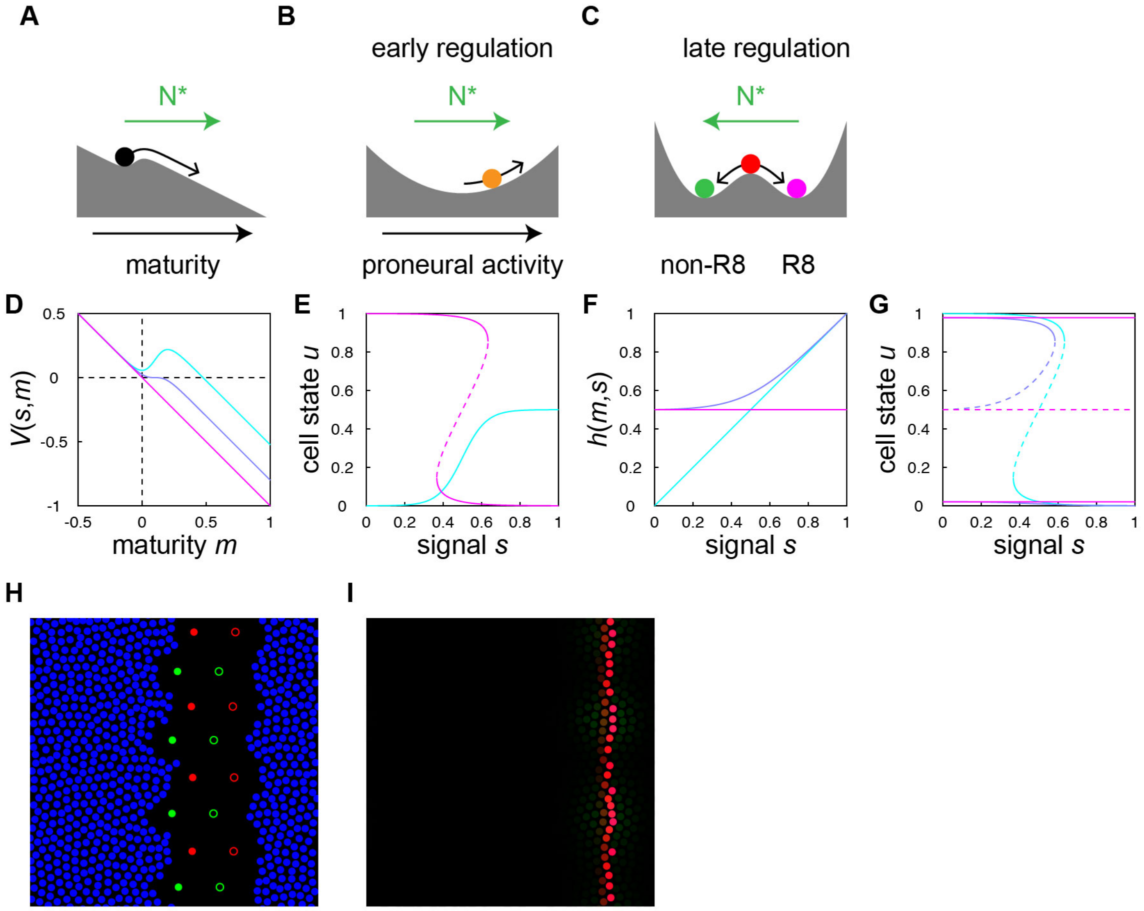

**Figure S10: Mathematical model for pattern propagation**

(**A**-**C**) Sketch of model in which Notch activation is required for initiation of cell maturation (A), and maturing cells transition from an early regulatory regime in which Notch promotes proneural activity (B) to a late, bistable regime in which Notch promotes a non-R8 fate (C).

(**D**) Landscape $V(s, m)$ for signal-dependent initiation of cell maturation (cf. Eqs. S11 and S13-S16). In the absence of signal (cyan), immature cells are trapped in a minimum at $m = 0$. For sufficiently large signal levels (the critical signal level is shown in blue and the limit of large signal levels in magenta), the local minimum disappears and cells begin to mature. In more mature cells, the slope of the landscape tends to a signal-independent limit, such that $m$ increases at a constant rate.

(**E**) Bifurcation diagram showing the steady states of the cell state $u$ vs. signal $s$ in the early (cyan) and late (magenta) regulatory regimes, in which signaling promotes (resp. inhibits) proneural activity (cf. Eqs. S12 and S17-S19).

(**F**-**G**) Incorporation of commitment in the model. Regulatory input $h$ (F; cf. Eqs. S25-S28) and the corresponding bifurcation diagrams (G) in uncommitted cells ($\xi_{\text{low}} = \xi_{\text{high}} = 1$), in a regime in which cells with low proneural activity are irresponsive to signaling ($\xi_{\text{low}} = 0$, $\xi_{\text{high}} = 1$), and a regime in which all cells are irresponsive ($\xi_{\text{low}} = \xi_{\text{high}} = 0$).

(**H**) Ordered template use to initialize model simulations (cf. section S8.5). Red and green solid circles show template R8 cells in rows -1 and 0, respectively, while immature cells are shown in blue. Cells in the black region, which are within signaling range from sources

representing R8 cells in rows -3 and -2 (open circles), are deemed to have already undergone differentiation and do not participate in the dynamics.
(**I**) Uniform template generated as described in section S8.6, showing proneural activity in a file of cells.

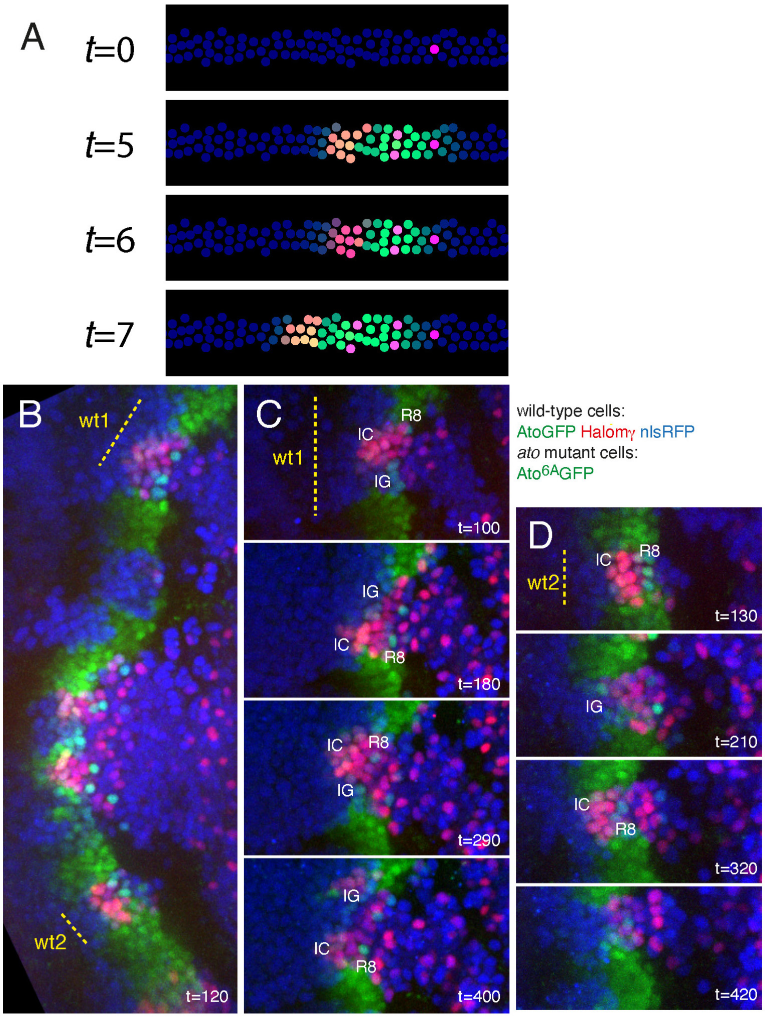

**Figure S12: Pattern propagation in one dimension**

(**A**) 1D pattern propagation in the model: simulation snapshots showing the quasi-1D propagation of the pattern in a strip of cells (labeled in blue).

(**B**-**D**) Live imaging of *ato*$^{6A}$*gfp* mosaic disc showing the dynamics of E(spl) (Halomγ, red) and wild-type Ato (AtoGFP, green) in wild-type cells (marked by nlsRFP, blue). Homozygous *ato*$^{6A}$*gfp* mutant cells (Ato$^{6A}$GFP, also green) are marked by the loss of nlsRFP (blue). These mutant cells do not carry Halomγ. The positions of two wild-type clones, wt1 and wt2 (dashed lines), which cross the differentiation front are shown (B). The temporal patterns of accumulation of Ato and E(spl) in the wt1 and wt2 clones are shown (C,D). New R8s and alternating teeth-forming pulses of Halomγ were detected every ~100 min in wild-type clones encompassing at least two columns (C), but only every ~200 min in wild-type clones encompassing a single column (D).

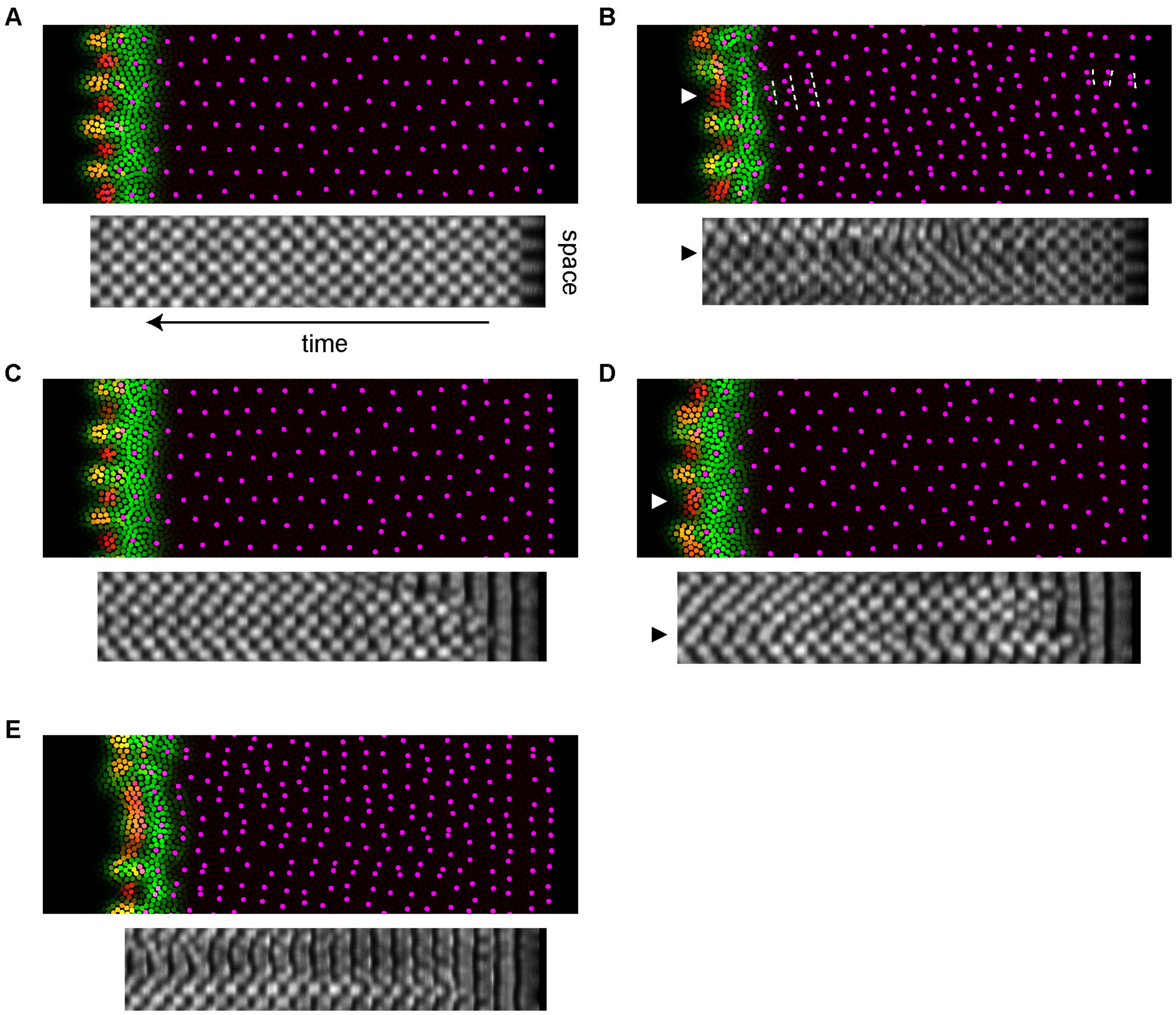

**Figure S13: Pattern propagation and symmetry breaking in the model**

(**A**) Simulated pattern initialized with a regular template and kymograph of the signal level at the leading edge of the pattern, illustrating the propagation of a regular pattern and accompanying signaling dynamics. The orientation and scale of the time axis are chosen such that regions of high signal level in the kymograph are approximately aligned with the R8s whose differentiation was initiated at that point in space and time.

(**B**) With a reduced basal signaling range, representing a *sca* mutant, orderly pattern propagation breaks down. Broader IGs can give rise to multiple R8s and these defects can repeat for several periods, as highlighted by dashed lines at some positions in the pattern. Arrowheads denote matching positions in space in the pattern and kymograph.

(**C**-**D**) Simulations with wild-type parameters and a uniform template. In C, a globally ordered pattern emerges, whereas in D, the dynamics is locally ordered but its phase is non-uniform in space, and a defect persists at the boundary between two ordered regions (arrowheads).

(**E**) Simulation with a reduced signaling range and a uniform template, showing a delayed symmetry breaking and an irregular pattern.

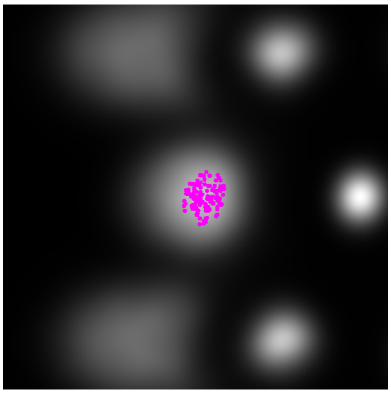

**Figure S14: R8 selection in the model**
R8 positions (n=128) from a model simulation overlaid on the average pattern of the cell state $u$ at the IG stage, showing that R8 cells emerge preferentially from the posterior side of IGs in the model, as in experiments (see Fig. S2J).

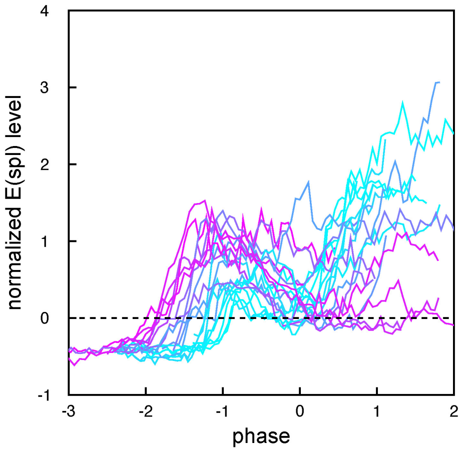

**Figure S15: Analysis of E(spl) dynamics in single cells**
Time courses showing the level of E(spl) in single cells (19 cells from 3 movies) as a function of the local phase of the E(spl) pattern, from the tracking of sparsely labeled nuclei. Tracks are colored according to the time at which the normalized E(spl) level first becomes positive, highlighting that a late onset of E(spl) expression at the IC stage (around phase -1; cyan tracks) correlates with an early re-expression of E(spl) at the IG stage (around phase 0).

**Table S1. Parameter values for initial model**

This table lists the parameter values used to simulate our initial model (Eqs. S3-S10, Fig. S1, movie S1). The signaling range $l$ is in units of the typical cell-cell distance $\lambda$, defined by $nl^2 = 1$, where $n$ is the density of cells per unit area.

| Description | Parameter values |
|---|---|
| Parameters taken from (*3*) | $l = 1.75$, $\tau = 1/2$, $a_0 = .05$, $a_1 = 1 - a_0$ |
| Parameters of the traveling front | $W = 16$, $c = 1$ |
| Template | $\Delta x = 3$, $\Delta \mathrm{y} = 8$ |

**Table S2. Parameter values for main model**

This table specifies the parameter values used for simulations of our main model (Eqs. S11-S29, Fig. 4E, Figs. S12A and S13A,C,D, movies S12, S13, and S16), as well as the parameter change used to simulate patterning in a *sca* mutant (Fig. S13B,E, movie S15).

| Description | Parameter values |
|---|---|
| wild-type | |
| Parameters taken from (*3*) | $l = 1.75$, $a_0 = .05$, $a_1 = 1 - a_0$ |
| Time scale for the dynamics of $u$ | $\tau = 0.3$ |
| Signal-dependent maturation | $k_s = 5$, $m_c = 0.2$ |
| Maturity-dependent dynamics of $u$ | $m_{\mathrm{early}} = 1$, $a_{\mathrm{early}} = 0.5$, $m_{\mathrm{late}} = 1.5$ |
| Maturity-dependent signaling | $m_{D^*} = 2$, $m_{l_{\mathrm{min}}} = 1.6$, $m_{l_{\mathrm{max}}} = 2.8$, $m_t = 3.1$ |
| Signaling teeth | $l_{\mathrm{min}} = X = 2$, $\Delta X = 3$, $l_{\mathrm{max}} = 7$, $w_t = 1.4$ |
| Commitment | $m_{\mathrm{low}} = 2.75$, $m_{\mathrm{high}} = 5$, $\tau_{\mathrm{high}}$=2 |
| Time scale for transitions | $\tau_{\mathrm{early}} = \tau_{\mathrm{late}} = \tau_t = \tau_{\mathrm{low}} = \tau_{D^*} = 2/3$ |
| Template | $\Delta x = 2.3$, $\Delta \mathrm{y} = 8$ |
| *sca* mutant | |
| reduced basal signaling range | $l = 1.25$ as in |

**Movie S1: Tentative model for pattern propagation by negative templating**
Simulation of our initial model for eye patterning, incorporating cell-intrinsic bi-stability and short-ranged inhibition, combined with a receding inhibitory front (panels and color code as in Fig. S1).

**Movie S2: Ato dynamics (1)**
Live imaging of AtoGFP (see Fig. 1D-D’’’) showing 5 pulses over a 8 hr movie (time in hr:min)

**Movie S3: Ato dynamics (2)**
Live imaging of AtoGFP (green; nuclear nlsRFP marker, red) showing traveling waves associated with the pulses of Ato which run from the equator to the poles (time in hr:min). AtoGFP expression was also detected in two groups of ocelli cells at the antero-dorsal margin of the eye disc (dorsal is up). Note that the ventral part of this eye disc was flipped over and that the AtoGFP signal was weak in this part presumably because most MF nuclei were outside the acquired volume.

**Movie S4: Ato dynamics in single cells**
Temporal dynamics of AtoGFP in single cells randomly labelled with nlsHalo (not shown). Three tracked cells are highlighted here to illustrate that cells in the MF can undergo 2 or 3 pulses prior to fate resolution (n=14 cells tracked from 2 movies).

**Movie S5: Average Ato dynamics**
Average dynamics of AtoGFP over three pulses (from IC stage to R8 stage; n=22 clusters from 2 movies; see Fig. S2I-I'''').

**Movie S6: Coordinated pulses of activity of the *ato3’* and *ato5’* enhancers**
Live imaging of ato3’-AtoGFP (green) and ato5’-AtoHalo (red) revealed coordinated pulses of activities in the MF (time in hr:min; see Fig. 2C,D).

**Movie S7: E(spl) dynamics**
Live imaging of GFPm$\delta$ and GFPm$\gamma$ revealed a periodic pattern of alternating teeth (time in hr:min; see Fig. 3A).

**Movie S8: Average E(spl) dynamics**
Average dynamics of GFPm$\delta$ GFPm$\gamma$ (see Fig. 3C), from n=15 positions in one movie, registered in space using PIV and in time according to the local phase of the dynamics.

**Movie S9: Dynamics of Ato and E(spl) reconstructed from fixed samples**
Animation of the average pattern of *ato* mRNA (green), *GFPm$\gamma$* mRNA (blue), and GFPm$\gamma$ protein (red) at successive phases in the dynamics (see Fig. S8A).

**Movie S10: Average dynamics of Ato and E(spl)**
Average dynamics of HaloAto and GFPm$\delta$ (see Fig. 3F), from n=16 positions in one movie, registered in space using PIV and in time according to the local phase of GFPm$\delta$.

**Movie S11: Landscape representation of dynamical model for R8 fate specification**
Cell differentiation trajectories in response to different signaling histories (upper panels) are depicted as the motion of balls in time-dependent landscapes. Whereas steady Notch signaling leads a non-R8 fate (left), low Notch activity following a differentiation-inducing pulse leads to an R8 fate (right). We note that this representation is intended to give an intuition of the model dynamics, rather than a fully accurate description, since the dynamics of the variables $\underline{m}$ and $u$ (Eqs. S11 and S12) do not strictly correspond to the gradient of a 2D potential.

**Movie S12: Simulation of pattern propagation in 2D**
Simulation of our main model, initialized with a regular template (see Fig. 4E; green, signal $s$; reg/magenta, cell state $u$).

**Movie S13: Simulation of pattern propagation in 1D**
Simulation of the model showing the quasi-1D propagation of the pattern in a strip of cells (see Fig. S12A).

**Movie S14: Dynamics of AtoGFP (green; nlsRFP, red) in *sca* mutants**
Live imaging of AtoGFP in *sca* mutant discs showed that patterning defects (lines of 2-4 R8s per column) propagate along the AP axis (time in hr:min; see Fig. 5E-E''').

**Movie S15: Simulation of the model with a uniform template and shorter-ranged inhibition**
Simulation of the model initialized with a uniform template, with a reduced basal range of signaling, showing a delayed symmetry breaking and an irregular pattern (see Fig. S13E).

**Movie S16: Simulation of the model initialized with a uniform template**
Simulation of the model initialized with a uniform template, showing the spontaneous emergence of an ordered pattern (see Fig. S13C).

**Movie S17: E(spl) dynamics in single cells**
Temporal dynamics of GFPm$\delta$ in single cells that were randomly labelled with nlsHalo. The intensity of the GFP signal in a single nlsHalo-positive nucleus that was tracked over time (left) is plotted (right; time in min).
